# Reward expectations direct learning and drive operant matching in *Drosophila*

**DOI:** 10.1101/2022.05.24.493252

**Authors:** Adithya E. Rajagopalan, Ran Darshan, Karen L. Hibbard, James E. Fitzgerald, Glenn C. Turner

## Abstract

Foraging animals must use decision-making strategies that dynamically adapt to the changing availability of rewards in the environment. A wide diversity of animals do this by distributing their choices in proportion to the rewards received from each option, Herrnstein’s operant matching law. Theoretical work suggests an elegant mechanistic explanation for this ubiquitous behavior, as operant matching follows automatically from simple synaptic plasticity rules acting within behaviorally relevant neural circuits. However, no past work has mapped operant matching onto plasticity mechanisms in the brain, leaving the biological relevance of the theory unclear. Here we discovered operant matching in *Drosophila* and showed that it requires synaptic plasticity that acts in the mushroom body and incorporates the expectation of reward. We began by developing a novel behavioral paradigm to measure choices from individual flies as they learn to associate odor cues with probabilistic rewards. We then built a model of the fly mushroom body to explain each fly’s sequential choice behavior using a family of biologically-realistic synaptic plasticity rules. As predicted by past theoretical work, we found that synaptic plasticity rules could explain fly matching behavior by incorporating stimulus expectations, reward expectations, or both. However, by optogenetically bypassing the representation of reward expectation, we abolished matching behavior and showed that the plasticity rule must specifically incorporate reward expectations. Altogether, these results reveal the first synaptic level mechanisms of operant matching and provide compelling evidence for the role of reward expectation signals in the fly brain.

## Introduction

An animal’s survival depends on its ability to adaptively forage between multiple potentially rewarding options(1, 2). To guide these foraging decisions appropriately, animals learn associations between options and rewards(3–5). Learning these associations in natural environments is complicated by the uncertainty of rewards, and both vertebrates and invertebrates employ decision-making strategies that account for this uncertainty (6–18). A commonly observed strategy across the animal kingdom is to divide choices between options in proportion to the rewards received from each(9–19). It has been hypothesized that animals that use this operant matching strategy make use of the expectation of reward – the recency-weighted rolling average over past rewards – to learn option-reward associations(18–20). Many studies further posit that this learning involves synaptic plasticity (21–23), and theoretical work has identified a characteristic relationship between operant matching and a specific form of expectation-based plasticity rule that incorporates the covariance between reward and neural activity(24–26). Despite this strong link between plasticity rules and the matching strategy, there has been no mapping of these rules onto particular synapses or plasticity mechanisms in any animal. As a result, deeply investigating these theories by manipulating and testing the nature of plasticity rules underlying operant matching has been intractable.

The fruit fly, *Drosophila melanogaster*, offers a promising system within which to address these challenges. Over the last half century, researchers have shown that flies can learn a wide variety of Pavlovian associations between cues and rewards(27–38). With the help of advances in functional and anatomical tools(39–45), they have identified the mushroom body (MB) as the neural substrate for these learning processes, including the assignment of value to sensory cues, and the underlying plasticity mechanisms have been extensively characterized(46–62). Recent theoretical work has also attempted to formalize the features of the learning rule that is mediated by these plasticity mechanisms(63–67). Despite this progress, evidence has been mixed as to whether this learning rule makes use of reward expectations(68–72), and there is a dearth of understanding about how flies learn in natural environments (but see (73, 74)). Studying foraging behaviors would allow us to not only clarify these gaps in the understanding of fly learning but could also provide an insightful framework for testing the neural computations underlying decision-making strategies such as matching.

Leveraging this foraging framework in flies requires us to address several open questions. First, animals in real foraging scenarios have to be able to form associations between multiple different options and rewards, yet evidence in flies suggests that some associations are labile and easily overwritten(34). Second, choice behavior has rarely been investigated at the individual fly level(74–76), and never in the context of flies making repeated choices between two probabilistically rewarding options. Whether flies can learn associations between options and probabilistic rewards and if so what behavioral strategies they use as a result, including whether they exhibit matching behavior, is therefore unknown. Finally, it is unknown whether they can integrate probabilistic reward events over multiple past experiences to form analog expectations. Even if such analog expectations can be formed, it is unclear if they lead to matching behavior through covariance-based plasticity in the fly brain.

We answered these questions using a novel olfactory two-alternative forced choice (2-AFC) task built for individual *Drosophila*. The assay allows us to measure hundreds of sequential choices from flies as we vary the probability of reward associated with two different odor cues. We found that flies can indeed learn multiple simultaneous associations between cues and probabilistic rewards, and this learning depended on the MB. We then designed a dynamic foraging paradigm inspired by mammalian tasks(14, 15, 18, 19) where reward probabilities associated with each odor change over blocks of many trials. We found that flies are able to keep track of these changing probabilities over time and adjust their behavior accordingly. We specifically showed that flies display operant matching behavior, and our analyses indicate that they do so by integrating information about reward and choice over multiple trials in the recent past. To test if this observed matching behavior requires particular types of plasticity rules we developed a MB-inspired model that generated de novo simulated behavior and found that matching was only observed if covariance-based plasticity rules that incorporated sensory expectation, reward expectation or both were utilized. Additionally, we restructured this model to predict fly behavior in our task and found that covariance-based rules better explained behavior than a simple rule that does not include expectation terms. We then directly demonstrated the requirement of a covariance-based plasticity rule that specifically incorporates reward expectation with experiments where we bypass computation of reward expectation by training animals with direct activation of dopaminergic neurons that drive plasticity. The behavior resulting from these bypass experiments were better explained by the simple plasticity rule that does not include expectation terms. Together these results offer the first synaptic level mapping and manipulation of learning rules underlying operant matching and provide compelling evidence for the role of reward expectation signals in the fly brain.

## Results

### Flies can learn multiple probabilistic cue-reward associations

We designed a Y-arena to study the decision-making strategies of individual flies when faced with probabilistically rewarding cues (Fig. 1A; Methods; Supp. Info. 1). The design of this assay was inspired by an earlier behavior assay for flies(77–81) and foraging related 2AFC tasks in vertebrates(13–15, 18). In our Y-arena, a single fly begins a trial in an arm filled with clean air and can choose between two odor cues that are randomly assigned to the other two arms (see Supp. Fig. 1A for estimates of odor boundaries). The fly can freely move between arms, with a choice defined as the fly crossing into the reward zone at the end of the arm (Fig. 1A). Once a choice is made, we provide reward by optogenetically activating sugar-sensing neurons using a Gr64f driver(82–84). The Y-arena then resets, with the arm chosen by the fly filled with clean air and the other two arms randomly filled with the two odors. This task design permits us to precisely control reward delivery without satiating the fly, and enables us to monitor the choices of a single fly over hundreds of trials.

**Figure 1:**
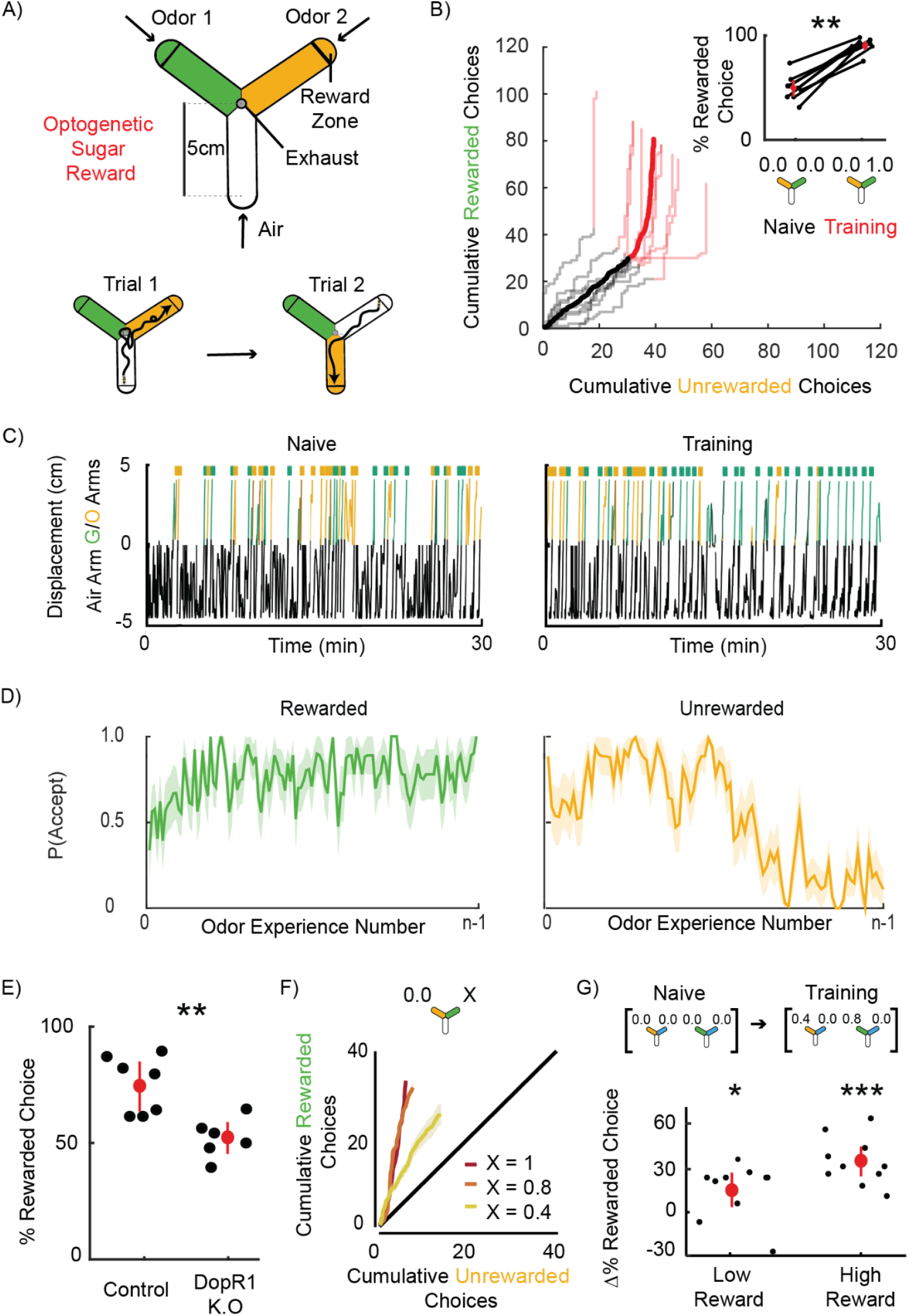
Flies learn multiple probabilistic cue-reward associations. **A** Schematic of Y-arena *(left)*. Airflow travels from tips of each arm to an outlet in the center. Reward zones are demarcated by lines (not visible to the fly). A choice is registered when a fly crosses into the reward zone of an odorized arm, triggering delivery of an optogenetic reward by activating Gr64f sugar sensory neurons with a 500 ms pulse of red light. The next trial then commences as the chosen arm switches to air and the two odors (represented as green and orange) are randomly reassigned to the other two arms *(right)*. **B** Cumulative choices made towards each option are shown (n = 9 flies, mean + individual flies). No rewards are available for the first 60 trials (Naive - black) and become available for the green option from the 61st trial onwards (Training - red). *Inset:* Percentage rewarded choices calculated naive (*left)* and training *(right)* blocks. Flies prefer the rewarded option in the training block compared to naive (Wilcoxon signed-rank test: p = 0.0039, n = 9). **C** An example trajectory of a fly in the Y before *(left)* and after *(right)* MCH (green) is paired with reward. Distance down the air arm is represented as negative values in black, while distances down an odorized arm are represented as positive values in green or orange to indicate odors. Choices are represented by colored raster ticks. At choice points the arena resets and that arm switches to air, so the fly’s position jumps to the tip of the air arm. **D** The probability of accept decisions are plotted as a function of time in the 100:0 protocol. Flies show a high probability of accepting the rewarded odor *(left)*. The probability of accepting the unrewarded odor is initially high and drops over time as rewards are made available *(right)*. **E** The percentage of choice made towards the rewarded option by control (*left*) and DopR1 K.O. (*right*) flies in one 100:0 block of 60 trials. Controls show a stronger preference for the rewarded option than DopR1 K.O. flies (Mann-Whitney rank-sum test: p = 0.0022). **F** Flies learn to associate odors with probabilistic rewards. Cumulative rewarded and unrewarded choices, across 40 trials, for three different reward probabilities : 1, 0.8, 0.4 *(top)*. Slope of all curves indicate that flies show a preference for the rewarded odor in all cases (Mann-Whitney rank-sum test: probability = 1, p = 4.4500*10^−8^, n = 18; probability= 0.8, p = 5.8927*10^−5^, n = 10; probability = 0.4, p = 0.0014, n = 10). **G** *Top:* Schematic of the protocol for training flies with two simultaneous probabilistic cue-reward contingencies. Two different odor choices are interleaved throughout a naive block of 80 total trials, and a reward block, where options were rewarded with probability of 0.4 or 0.8. The blue arm indicates that the unrewarded odor is always the same. *Bottom:* Performance (percentage of choices in which potentially rewarding option was chosen) on the low and high reward choices, showing that individual flies learn both associations (Mann-Whitney rank-sum test: p = 2.3059*10^−4^ for high rewarding odor; n = 10 p = 0.01 for low rewarding odor, n = 10).

We first established that flies learn effectively in this apparatus by reliably rewarding flies only when they chose one of the odors - what we term a 100:0 protocol. Each fly first experienced the two odors (3-octanol; OCT and 4-methylcyclohexanol; MCH) unrewarded for a block of 60 trials, and then reward delivery was activated for the following block of 60 trials. As observed previously, although individual flies exhibited different odor biases in this naive phase(75, 76, 85), those biases averaged out over the population (Fig. 1B + inset). In this phase, Flies spent a lot of time in the air arm and made variable choices, with little preference for either odor (Fig. 1C left, example fly). Once reward was made available, flies rapidly shifted to choosing the rewarded odor (Fig 1B). This was accompanied by a faster interval between choices (Supp. Fig. 1B), and a decrease in meandering trajectories (Fig. 1C).

To analyze this choice behavior at a more elemental level, we adopted the common framework of considering foraging choices as a series of accept-reject decisions, where the animal decides whether or not to pursue an option(86). We defined reject decisions as when a fly enters an odorized arm but turns around and exits the arm before reaching the reward zone, while accept decisions reflect cases where the fly reaches the reward zone (see Methods). Associating options with rewards changed the probability of accept decisions gradually over the course of a block. Acceptance probability increased for the rewarded odor and decreased for the unrewarded odor (Fig. 1D, Supp. Fig. 1E). On average, flies were around four times more likely to reject the unrewarded odor and seven times more likely to accept the rewarded odor (Supp. Fig. 1D). Interestingly, flies tended to reject odors quite close to the tip of the arm (Supp. Fig. 1F), suggesting that flies might accumulate evidence over time to make and commit to their decision – an aspect of fly behavior that has previously been studied(87). These results indicate that fly choice behavior in this task can be thought of as a series of accept-reject decisions.

We found that the odor-reward associations learnt by flies in our assay were MB dependent. Learning-related plasticity in the MB circuit requires the activity of dopaminergic neurons (DANs) (33, 34, 51, 55–58, 60, 88). Dopamine is sensed by odor-representing Kenyon cells (KCs) and induces synaptic plasticity between these KCs and downstream mushroom body output neurons (MBONs)(33, 55, 57). To interfere with this plasticity, we used a tissue-specific CRISPR knock-out strategy(89) to knock out DopR1 receptors selectively in the KCs (Methods), which are necessary for flies to associate odors with rewards in other paradigms(60). These flies showed no detectable learning in the 100:0 protocol, compared to control animals (Fig. 1E). These findings establish that odor-reward associations in our novel behavioral essay are mediated by MB plasticity.

We then asked whether flies could link odor cues with probabilistic rewards and distinguish between different reward probabilities, a key aspect of natural foraging. Importantly, we incorporated reward baiting into our probabilistic reward tasks(15, 18). This means that rewards probabilistically become available and then persist until the reward is collected (Methods). Baiting is commonly used in mammalian 2AFC tasks, as it is thought to mimic the natural processes of resource depletion and replenishment over time. We began with experiments in which a single odor was rewarded with a range of probabilities: 1, 0.8 or 0.4. Flies showed a preference towards the rewarded odor in all cases (Fig. 1F). The extent of the preference varied with the probability of reward - a higher probability of reward led to a stronger preference. Interestingly, flies made faster choices when rewards were more probable (Supp. Fig. 1C).

These results show that flies can learn from probabilistic rewards but do not determine if they can store two associations simultaneously - another necessity for foraging. To test this, we designed a paradigm with a third odor, pentyl acetate (PA) included. This served as the unrewarded cue while we tested memory formation with the other two odors (Fig. 1G top). Flies first made 80 unrewarded choices consisting of 40 choices between OCT and PA and 40 choices between MCH and PA. In the next 80 (Training) trials, one of OCT or MCH was assigned a high reward probability (0.8) and the other a low probability (0.4). We interleaved the training trials for the two different odors, to ensure both relationships would be learnt simultaneously. After pairing, flies preferred both rewarded odors over PA (Fig. 1G bottom). This choice preference was also reflected in their accept/reject behavior, with flies exhibiting a clear preference for accepting the high-rewarding odor (Supp. Fig 1G right). Interestingly, in trials with the low-reward cue presented, there was an increased probability of rejecting both rewarded and unrewarded odors, as compared to naive trials (Supp. Fig. 1G left). This suggests the possibility that flies keep track of all the odor options potentially available in the environment, and actually increase their rejection rate in the absence of the high-reward odor.

Overall, these experiments establish the fly as a capable animal model for studying foraging behaviors. Individual flies in the Y-arena can learn multiple odor-reward associations and can do so in the face of probabilistic reward. Importantly, these relationships are mediated by synaptic plasticity at the KC-MBON synapses in the MB. This establishes a foundation to test how these animals perform in dynamic foraging tasks, and assess how they respond to reward probabilities that change over time.

### Flies follow Herrnstein’s operant matching law

Foraging tasks are cognitively complex, involving two cues paired with different reward probabilities that change with time. This requires animals to keep track of choice and reward history and form expectations to make adaptive choices. We designed our own dynamic foraging protocol to investigate how flies behave in such a scenario. The protocol consisted of three consecutive blocks of 80 trials each. Flies made choices between two odors (OCT & MCH) that were paired with different reward probabilities (Methods). These probabilities remained fixed within a block and changed across blocks (Fig. 2A, example).

**Figure 2:**
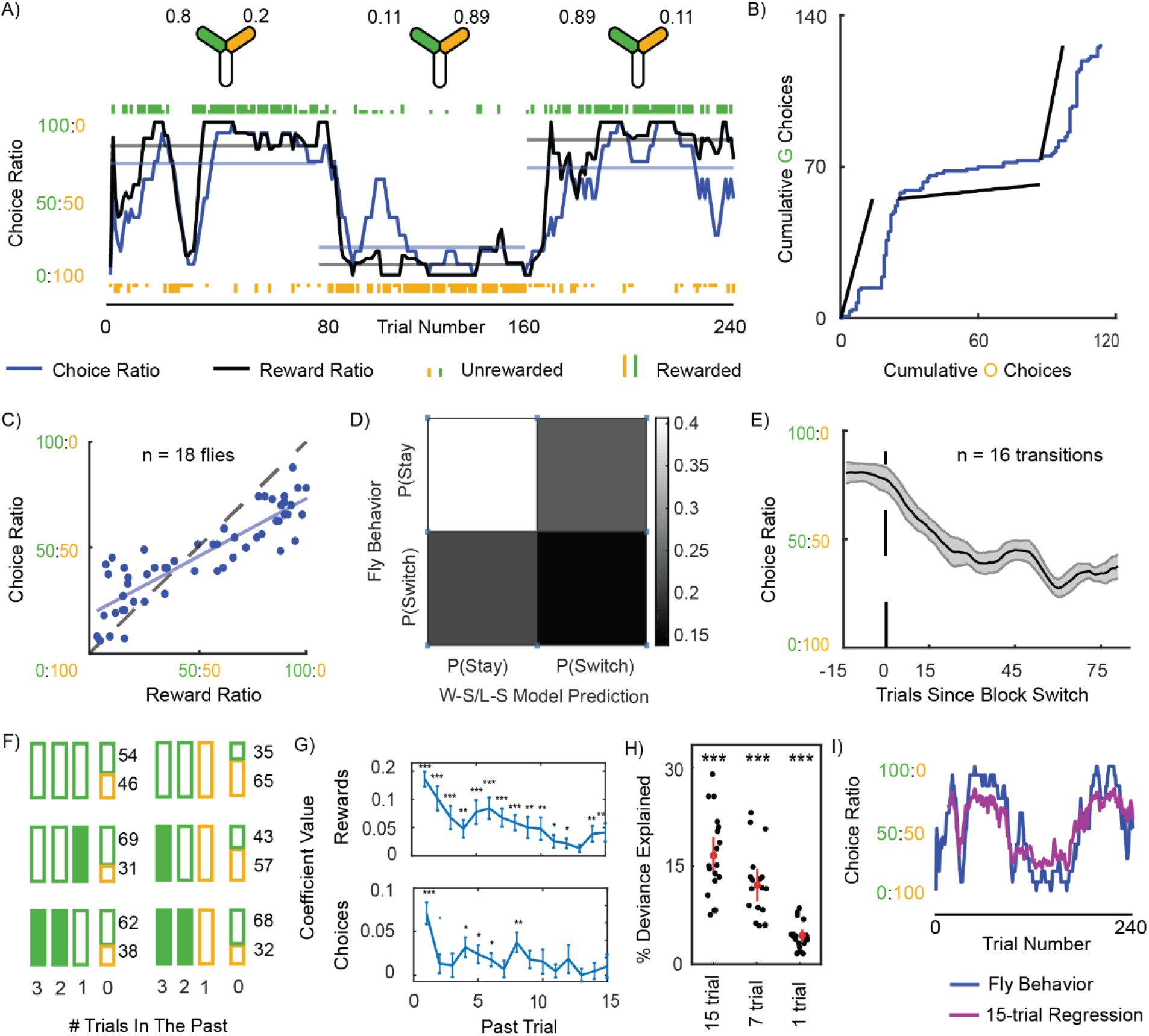
Flies follow the operant matching law. **A** Matching of instantaneous choice ratio (blue) and reward ratio (black) in an example fly. Reward probabilities for trial blocks are indicated by the Y-arena icons above *(top)*. Individual odor choices are denoted by rasters, tall rasters - rewarded choices, short rasters - unrewarded choices. Curves show 10-trial averaged choice ratio and reward ratio, and horizontal lines the corresponding averages over the 80-trial blocks. **B** Cumulative choices of the same example fly. The slope of the black lines indicate the block-averaged reward ratio in the three successive blocks; the blue line indicates the cumulative choices with slope representing choice ratio. The parallel slopes of the two lines indicate matching. **C** Block-averaged choice ratio is approximately equal to reward ratio, following the matching law, but with some undermatching (n=54 blocks from n=18 flies). **D** Confusion matrix indicating the probability of true and false switches and stays predicted by the “win-stay; lose-switch” model when compared to fly data. **E** Change in instantaneous choice ratio around block changes (n = 16 transitions with large changes in reward probabilities between blocks). **F** Analysis of choices following particular histories of experience. Choices made by flies over three consecutive past trials are represented by boxes of different colors. Colors represent odors chosen, and rewarded choices are represented by filled boxes. Probabilities of choosing the green and orange odor on the current trial conditional on this history are illustrated with associated values. 6 out of 64 possible sequences are illustrated here. **G** Coefficients from logistic regression performed on fly choice behavior to determine the influence of 15 past rewards *(top)* and choices *(bottom)* on a fly’s present choice (Mann-Whitney rank-sum test: *** - p < 0.001, ** - p < 0.01, * - p<0.05, n = 18 flies). **H** Model fit quality (percentage deviance explained) for 15-trial logistic regression, 7-trial logistic regression and 1-trial logistic regression models. Null model used to calculate the quality metric is a logistic regression with 0-trial history and only bias. **I** 15-trial logistic regression fit (purple) on behavior (blue) from the example fly from panel A, plotted from the 15th trial onwards to avoid edge effects.

We found that flies exhibit operant matching behavior, similar to observations in monkeys, mice and honeybees(11, 14, 15, 18). Individual flies exhibited a strong correlation between choice ratio and reward ratio either calculated over entire blocks or over a short (ten-trial) window, to capture short-term dynamics/fluctuations (Fig 2A,B - example fly; Supp. Fig. 2 - all 18 flies). This holds true across flies, as seen in the relationship between block-averaged reward ratios and their choice ratios (Fig. 2C). In such a plot the matching law predicts that all points will fall along a line with slope equal to one (the unity line). Flies appear to approximately follow the matching law with a slight amount of undermatching, signified by a slope less than one. Undermatching is commonly observed across species(14, 15, 17–19), and several reasons have been suggested for this tendency(17, 26) (see Discussion).

Past work has suggested that animals form expectations of reward and use this to guide behavior in such dynamic foraging tasks(17–19, 24). When rewards are delivered probabilistically, animals can only derive an expectation of reward by tallying information over multiple trials. However, such tallying could reflect a computation beyond the capabilities of flies. We wanted to explicitly address the alternative hypothesis that flies follow a simple win-stay/lose-switch strategy (Supp. Fig. 3A), which would suggest that their behavior is dictated by only the most recent reward/omission experience. Simulating choice sequences using this learning rule produced output that qualitatively resembled that of the fly (example in Supp. Fig. 3B). However, it poorly captured the stay/switch probabilities actually observed in fly behavior data (Fig 2D). In particular, switching occurred much more frequently than predicted. As further evidence that multiple past outcomes affected behavior, choices of an individual fly at block transitions showed a lag between the choice ratio curve and the updated reward ratio at transition points (Fig. 2B), suggesting that the fly takes a few trials to adjust its behavior. Quantifying this across multiple transitions for all flies in the task showed flies require 15-20 trials to reach a new steady state choice behavior following block switches (Fig. 2E). To test the role of multiple past trials on present choice more explicitly, we first looked at the decision made by flies on a given choice as a function of their past choices and outcomes. We focussed on choices following particular triplets of past experiences, inspired by recent work in mice(8) (Fig. 2F). For example, following three unrewarded choices of one particular odor, flies’ next choice was roughly random. However, when an odor was rewarded on the most recent trial or more distant trials, choices were biased towards that option (Fig 2F left). In another comparison, flies’ tendency to switch back to an earlier choice (i.e. choose the green odor after an unrewarded choice of the orange odor) increased based on how that odor was rewarded in the past (Fig 2F right).

**Figure 3:**
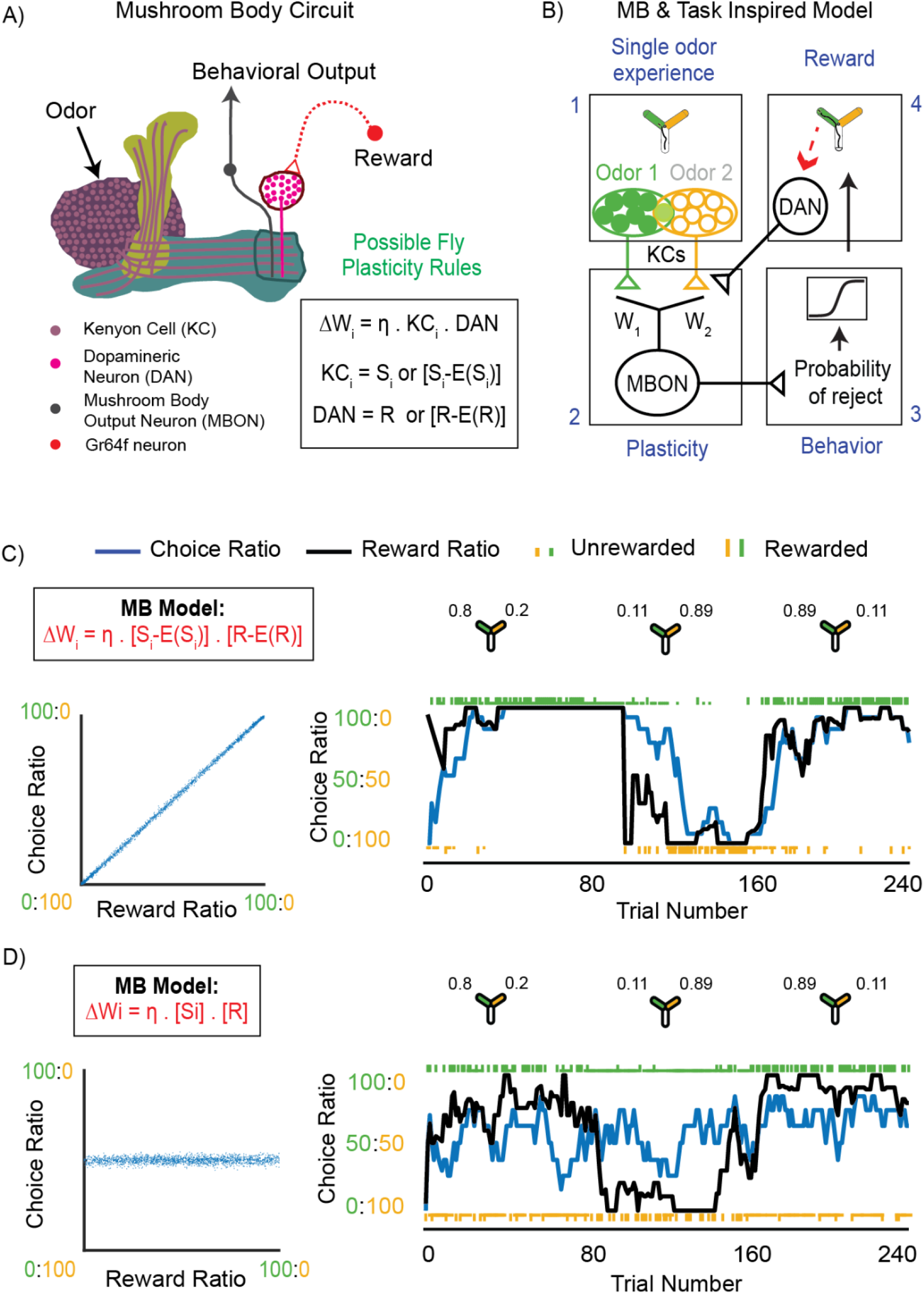
Covariance-based learning is required for matching behavior in a model of the MB. **A** Schematic representing the MB with all relevant neurons shown in different colors (*key*). *Box - right:* reward dependent synaptic plasticity rule at the KC-MBON synapse. **B** Schematic adapting the model developed by L&S to more closely resemble the MB and the features of our olfactory task. *1*. Sensory inputs represented by populations of KCs with overlap between representations. In the modified task, agents only experience one odor at a time. *2*. Weights between inputs and MBON are modified according to plasticity rules shown in A. *3*. MBON output determines probability of rejecting an odor and is passed through a sigmoidal nonlinearity to determine behavior. *4*. Reward information is provided to this circuit via DAN activity which either represents simply reward (R) or reward minus reward expectation (R-E[R]). **C** *Left:* Block-averaged choice ratio produced by the [S_i_-E(S_i_)]*[R-E(R)] covariances-based rule (*box*) plotted against reward ratio. The model exhibits matching behavior (slope is 1). *Right:* An example simulation showing the performance in a 3 block task of a model incorporating a covariance-based rule [S_i_-E(S_i_)]*[R-E(R)]. Task reward contingencies are the same as shown for the example fly in Fig. 2A. **D** Same as (C), but simulated with a non-covariance learning rule. *Left:* The model produces behavior that does not show matching (slope < 1). *Right:* performance in a 3 block task does not show matching of choice and reward ratio.

To measure the relationship between current choice and past outcomes more systematically, we used logistic regression to determine how a fly’s decisions depended on choice and reward history. Like other animals(8, 15), flies showed a small amount of habitualness choosing options that had been recently chosen more often; regression coefficients for a small history of most recent choices were significantly positive (Fig. 2G bottom). However, more significantly, this approach showed that the reward history was important for predicting choice, with recent rewards weighted more than those in the more distant past; regression coefficients for the 15 most recent rewards were significantly non-zero across the population of flies (Fig. 2G top). Further, we compared regression models that predicted behavior based on different lengths of outcome histories (15, 7 and 1 trial) and found that the percentage of deviance explained over a null model with a 0-trial history was great for models that used longer outcome histories (Fig. 2H; Methods). An example fit from a regression model with a 15-trial history is shown(Fig. 2I). Additionally, we fit behavior of individual flies with a leaky integrator model(18) which assigns value to options using exponentially weighted reward histories (Supp. Fig. 3C-G). This analysis found that an average exponential timescale of 7 trials was best for predicting behavior (Supp. Fig. 3D), in agreement with the regression coefficient of rewards 7 trials in the past being about half the value of the coefficient of the most recent reward in the logistic regression model (Fig. 2G). Both these approaches support the finding that flies integrate several past trials and don’t rely on just the most recently experienced outcome. Together, these results show that flies’ choices follow operant matching, with each choice depending on the history of many past outcomes.

### Covariance-based learning is required for matching behavior in a model of the MB

Theoretical work has placed strong, testable constraints on the nature of learning rules that could underlie this ubiquitous operant matching strategy. An elegant theory put forward by Loewenstein and Seung(24) proves that operant matching is the inevitable outcome of synaptic plasticity rules that modify value-representing synaptic weights according to the covariation of reward and neural activity. Here covariance is defined by the product of these two variables, with at least one of these terms being subtracted by its expectation or mean (Supp. Fig. 4A - box). The steady-state behavior of covariance-based rules provides an intuition for their link to operant matching(24). For example, consider a rule that only subtracts the reward term by its reward-expectation(Supp. Fig. 4A - box). If such a rule is at a steady state when animals forage, which means that synapses on average are not changing anymore, it will require the animals to make choices such that the rewards they receive are equal to the reward expectation. This requirement nicely resonates with the definition of operant matching. By definition, operant matching is a strategy in which choices are divided between options such that the average number of rewards received over a given number of trials is the same for both options. Only an animal that follows operant matching would receive rewards at a rate equal to the reward expectation for both options, suggesting that the steady state behavior of a covariance-based rule has to be operant matching. Loewenstein and Seung mathematically formalized this intuition and further showed that operant matching can only arise from covariance-based rules when synaptic plasticity is involved.

Loewenstein & Seung used a simple neural circuit model to illustrate their theory (Supp. Fig. 4A). In this toy-model choice options were signaled by two sensory neurons, which synapse onto two motor neurons to drive two different actions. The value associated with each option is represented in the weight of those synapses, and the outcome of a decision is based on a winner-take-all interaction between the motor neurons - whichever is strongest dictates the decision. We simulated this model circuit and confirmed that operant matching only arises when synaptic weights are updated according to covariance-based rules (Supp. Fig. 4B-E). Interestingly, the structure of this model maps nicely onto the circuitry of the fly MB (Fig. 3A). The odor choices are represented by the KCs, each odor activating a sparse subset of the KC population(90–93). KCs synapse onto MBONs, which guide action by signaling the valence of an odor i.e. it’s attractive/repulsive quality, rather than a specific action(31, 56, 62). KC-MBON synapses are modified by a plasticity rule that depends on the coincident activity of odor-representing KCs and release of dopamine by reward-signaling DANs (33, 34, 51, 55–58, 60, 88, 94) (Fig. 3A - box). Current evidence indicates that post-synaptic activity of the MBON does not play a role in the plasticity (47), so only the covariance between sensory and reward activity needs to be considered.

This overall structure aligns reasonably well with the toy-model, however there are differences both at the level of the MB circuitry and the structure of our task. While the theory of L&S is incredibly general and makes very limited assumptions about the nature of any potential underlying neural circuitry, we wanted to ensure that incorporating the biological realities of MB circuit architecture does not affect the relationship between covariance-based rules and matching. First, rather than having two input neurons, odors are represented by noisy populations of KCs. We thus parameterized input representation in the model to incorporate noise and overlap of KC subsets between options (Fig. 3B-1; Methods). Further, flies only experience one odor at a time in our task. Therefore only one of the inputs (and resulting overlap in the other input) to the model agent were active at any time (Fig. 3B-1, 3B-4). Secondly, plasticity between MBONs and KCs are modified by a synaptic depression rule(55, 56). We thus flipped the sign of the weight update rule to produce depression from positive covariance signals (Fig. 3B-2). Finally, output in the MB is driven by the MBONs, whose activities determine whether flies accept or reject an odor rather than a winner-take-all choice mechanism(30, 31, 56) (Fig. 3A). We modeled output to reflect this by having model MBON activity encode the probability of rejecting an odor, with higher activity having a greater probability to reject. To account for possible stochasticity in this process, we passed model MBON activations through a sigmoidal nonlinearity to determine behavioral output (Fig. 3B-3).

With this modified model in place we simulated behavior resulting from covariance rules that incorporated stimulus-expectation, reward-expectation or both. All three covariance-based rules that produced matching in the original model continued to produce matching in our modified model (Fig. 3C left; Supp Fig. 5A,B left), consistent with the theory. We also observed that the covariance-based rules could replicate the trial-by-trial behavior of flies in the dynamic foraging paradigm, tracking changes in the reward contingencies across blocks, with the resulting instantaneous choice ratio biased towards the more rewarded option in each block (Fig. 3C right; Supp Fig. 5A,B right). In contrast, a rule that did not incorporate either reward or stimulus expectation did not produce matching (Fig. 3D left) and did not appropriately capture instantaneous behavior either, with choices made roughly equally to both options throughout (Fig. 3D right). This deviation from matching is in agreement with Loewenstein and Seung’s theory. It also makes intuitive sense if one considers the nature of this non-covariance rule. A rule that depends simply on the product of neural activity and the presence of reward will only cause unidirectional plasticity, eventually pushing synapses related to both options to saturation though at slightly different time-scales. This would result in both options being chosen roughly equally. These results, paired with our observation of operant matching in *Drosophila*, indicate that flies use a covariance-based rule to learn option-reward associations in the MB.

### Identifying learning rules underlying dynamic foraging in the mushroom body

To test if our theoretical prediction of a covariance-based rule is supported by the observed behavior, we developed an approach that estimated the form of the plasticity rule being used in the fly MB. Our goal was to break the plasticity rule into components that span a large space of possible rules in the fly MB and use optimization approaches focussed on accurately predicting behavior to assign optimal coefficients to each of these components. In this way we would identify the form of the plasticity rule that best explained observed behavior and be able to conclude if this rule was a covariance-based rule.

The approach we developed made use of the structure of the MB-inspired generative model described in the previous section (Fig. 3B) to predict accept/reject decisions made by the fly based on sensory and reward inputs (Fig. 4A, Methods). The key distinction here was that rather than incorporating a pre-defined plasticity rule such as one of the covariance-based rules, this model used a rule that consisted of four terms that spanned the space of biologically-plausible rules in the MB. We then transformed this into a logistic regression problem (Methods). At every iteration of the regression, behavior predicted by the model was compared to true fly behavior and regression coefficients assigned to each component of the plasticity rule was updated. The set of coefficients for each of the components of the plasticity rule that minimized the error between predicted behavior and the fly’s actual behavior could then be identified as the plasticity rule used in the fly MB.

**Figure 4:**
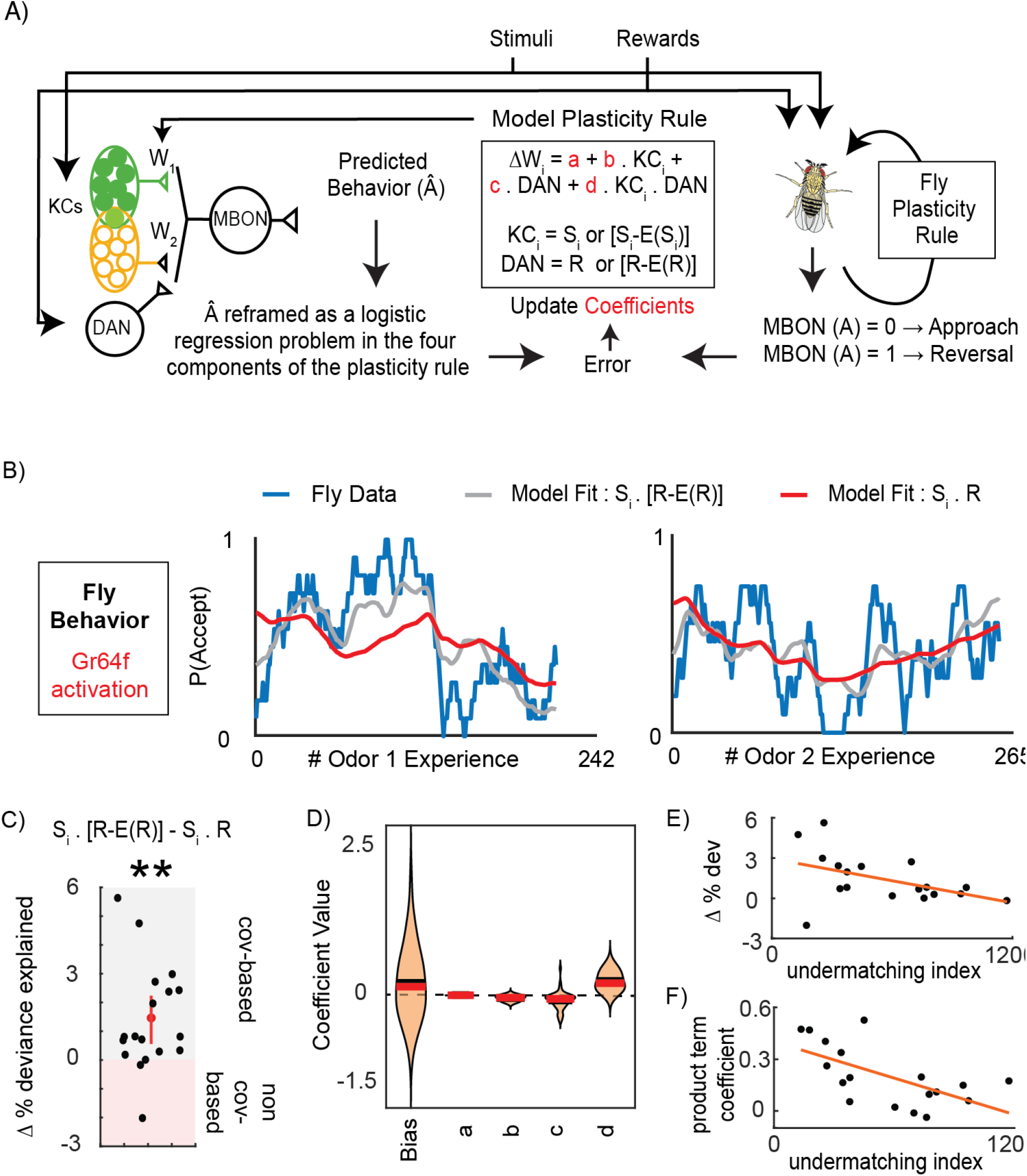
Identifying learning rules underlying dynamic foraging in the mushroom body. **A** Schematic detailing the logic of the MB-inspired regression model. **B** Example fly data (blue) showing the probability of accepting odor 1 *(left)* and odor 2 *(right)* calculated over a 6-trial window as a function of the number of times the fly experienced the given odor. This data was fit using an MB-inspired regression model (A) that incorporates either an covariance-based rule with reward-expectation (gray), or a non-covariance rule (red). **C** Change in percentage deviance explained, computed by subtracting the percentage deviance explained of the non-covariance-based model from a covariance-based rule that incorporates reward expectation (n = 18 flies). On average, fly behavior was better predicted by the covariance-based model (Wilcoxon signed-rank test: p = 0.0018). Individual flies that were better fit by the covariance-based model have a positive value on this plot (gray region) while flies better fit by the non covariance-based model have a negative value (red region). **D** Regression coefficients assigned to each term of the plasticity rule when the MB-inspired regression model using a covariance-based rule with reward-expectation was fit to the flies’ behavior. Largest coefficients were assigned to the product term. **E** Change in percentage deviance explained (shown in C), plotted against a measure of undermatching (mean square error between instantaneous choice ratio and reward ratio lines) for each fly (n = 18). **F** Coefficient value assigned to the product term term (shown in D), plotted against a measure of undermatching for each fly (n = 18).

The activity of KCs and DANs are the only known factors to affect synaptic plasticity(47, 63). We thus included in the model four terms: a constant term, a KC-related term, a DAN-dependent term, and a term dependent on the product of KC and DAN. (Fig. 4A center box). KC terms reflect stimulus information while DAN terms reflect reward information. We set up the model so that these terms may or may not be subtracted by their respective expectation values. By definition, when either of these terms incorporates the mean subtraction, the DAN, KC product term becomes a covariance term. This resulted in four different models, three of which could give rise to covariance-based rules if the product term was assigned a large coefficient. This method allowed us to explore the space of possible plasticity rules in an unbiased way and ask if covariance-based rules were in fact the rules that best predicted fly behavior.

We first validated this approach by simulating choice sequences using each of the three covariance-based rules, as well as the simple non-covariance rule, and checking whether the model fitting identified the correct rule. Indeed, data simulated with a covariance-based rule that included reward expectation was better fit by a model that incorporated this reward expectation subtraction into its plasticity rule (Supp. Fig. 6A,B). Moreover, the model correctly assigned the largest weights to the DAN-KC product term, the term that calculates covariance between these two elements(Supp. Fig. 6C). We also found that the extent of matching produced by the model was largely unaffected by the degree of overlap in KC activity patterns (Supp. Fig. 6D-G), and the regression model accurately recaptured simulated data when biologically-plausible overlap values(92) were used (Supp. Fig. 6H). Similar analysis revealed that the approach also gave consistent results across a range of timescales for calculating accrued value(Supp. Fig. 6I). We therefore assumed that overlap was 0 and used an exponential timescale of 3.5 trials (similar to logistic regression in Fig. 2G) in all future analysis.

We then applied our approach to fit data from flies performing the dynamic foraging protocol. Regressions that utilized expectation subtractions in their plasticity rules usually captured fly behavior better, as can be seen with a representative example fly (Fig. 4B, Supp. Fig. 7B). To compare fit quality of the different models for every individual fly, we calculated the percentage deviance explained for each. This metric showed that rules that subtract sensory expectation, reward expectation or both were better fits for fly behavior (Fig. 4C, Supp. Fig 7A). Interestingly, we found that in some flies the simple expectation free non-covariance rule was a better fit. One explanation for this result is that these flies showed operant matching to a lesser extent. We quantified matching by calculating the mean squared error between instantaneous choice and reward ratios. This analysis found that flies did exhibit different strengths of matching and that this was correlated with how well an expectation free plasticity rule fit the behavior data. Flies that were better fit by the expectation free rule tended to show more undermatching, in line with our predictions (Fig. 4E).

Regressions that subtracted sensory, reward or both expectations were generally better fits for fly behavior. However, this alone does not mean that a covariance-rule is used by the fly. This would only be true if the largest coefficient that results from these regressions was assigned to the DAN-KC product term. We therefore examined the regression coefficients resulting from this analysis to assess which terms were important for predicting behavior. When a rule that incorporated reward expectation was used to fit fly behavior, we found that the best-fit model parameters had a similar profile as when this rule was used to fit simulated data in Supp. Fig. 6A. The largest regression coefficients were assigned to DAN-KC product term showing that flies do in fact use a covariance based rule to guide their behavior (Fig. 4D). We compared the value assigned to the product term for a given fly against the extent of matching and found this coefficient was larger for flies that matched better (Fig. 4F), further clarifying that this product term in the plasticity rule is related to the extent of matching, as expected by the Loewenstein and Seung theory. Other parameters were on average assigned values near zero with some fly-to-fly variability (Fig. 4D; Supp. Fig. 7C,E,G) and correlations between components that were typical of statistical models yet hard to interpret (Supp. Fig. 7D,F,H,I). This importance of the product term was also observed when sensory expectation or both sensory and reward expectation were incorporated into the plasticity rule (Supp. Fig. 7C,E,G). Together, these results link the previous theoretical finding to real behavioral data and further support the idea that a covariance-based rule is implemented in the MB.

### Behavioral evidence of reward expectation in DANs

While the regression analysis confirmed that covariance-based rules were better at predicting fly behavior, it could not distinguish between different covariance-based rules. However the mathematical differences between the three rules suggested a way forward. In particular, the rules differ in which terms - sensory input or reward - incorporate an expectation. Thus, to distinguish between the possible different covariance-based rules in the MB, we designed an experiment to manipulate the calculations of reward expectation using genetic tools that override the natural activity of the DANs. Specifically, we provided reward via optogenetic activation of the reward-related protocerebral anterior medial (PAM) DANs. This would bypass any upstream computation of reward expectation and simply provide a consistent reward signal on every trial. Such a manipulation would change the learning rule from a covariance-based rule to a non-covariance rule if the following conditions were met: i) the animal’s learning rule depended on the product of DAN and KC activities; ii) DAN activity incorporated reward expectation; iii) KC activity did not incorporate sensory expectation. This would in turn result in modified behavior. For this test we initially focused on a task consisting of two blocks (naive and training) of 60 trials each, with a reward ratio of either 100:0 or 80:20.

We first predicted how the behavior in these protocols would differ between covariance-based and non-covariance rules using simulations. As expected, covariance-based models learnt to choose the more rewarded option more often, with choice ratios reflecting reward probabilities (Fig. 5A, Supp. Fig. 8A,B). On the other hand, non-covariance rules failed to show a strong effect on the choice ratio, and preference saturated around 75% in 100:0 and 50% with the 80:20 reward ratio (Fig. 5B). These theoretically predicted preferences very closely match our observations of fly behavior in the DAN bypass experiments. We observed low plateau performance in both tasks (Fig. 5F), with values strikingly similar to that predicted by the non-covariance rule (Fig. 5D). The behavior of the model with any covariance-based rule was similar to the fly behavior when it was rewarded using the sugar neurons (Fig. 5C,E).

**Figure 5:**
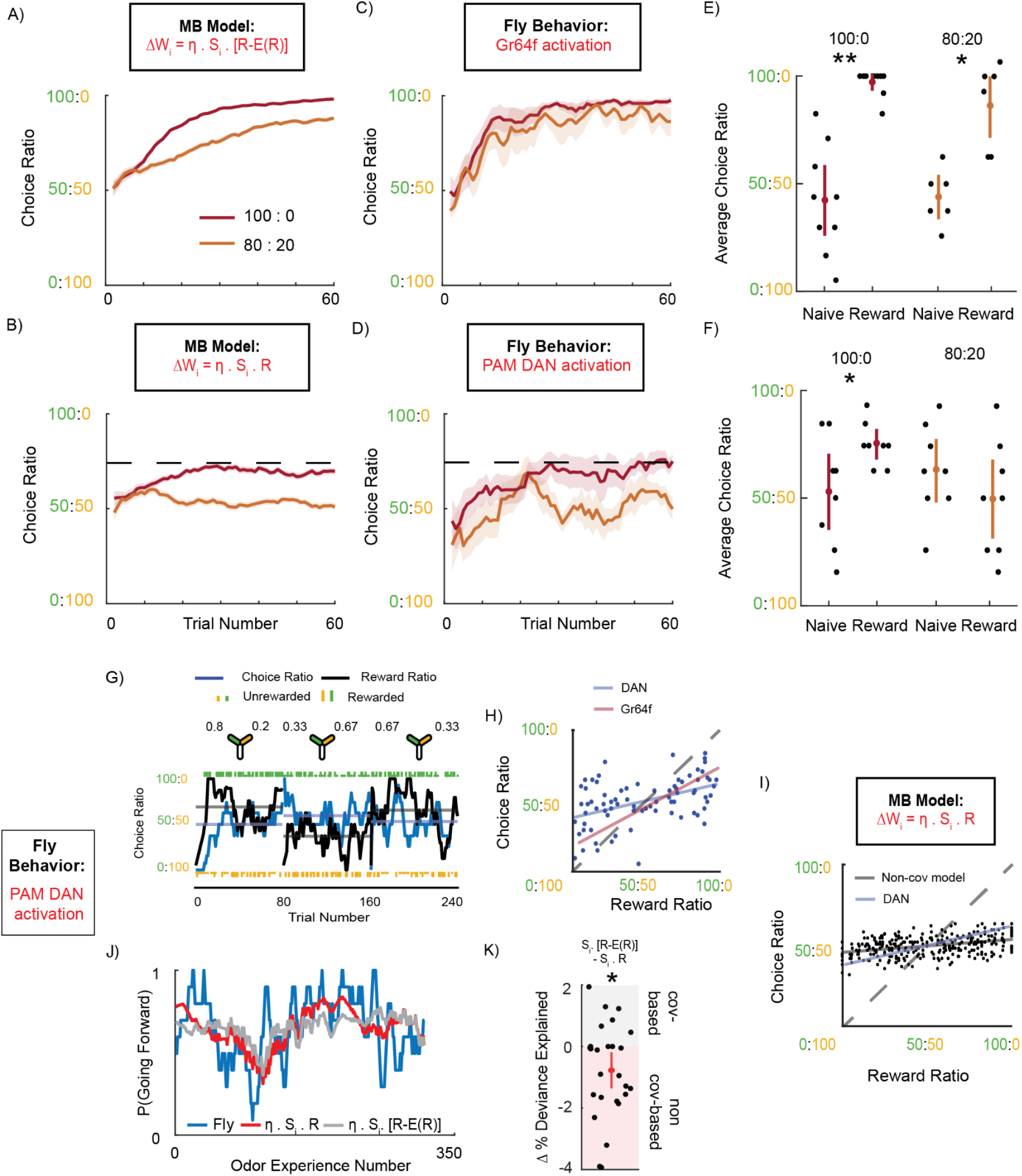
Behavioral evidence of reward expectation in DANs. **A** Simulated instantaneous choice ratio plotted as a function of trial number, for an agent using a covariance-based rule with reward expectation in 80:20 (orange) and 100:0 (red) reward conditions. **B** As (A), except for an agent using a non covariance rule. Dashed line indicates the maximum possible performance of this agent in the 100:0 protocol. **C** Flies’ instantaneous choice ratio when providing optogenetic reward using the sugar sensing neuron driver Gr64F-Gal4 to drive UAS CSChrimson. Again reward ratios are 80:20 (n = 6 flies) and 100:0 (n = 9 flies). **D** As in C except reward provided via the PAM DANs using R58E02-Gal4 to drive UAS-CSChrimson (n=8 flies in both 80:20 and 100:0). Dashed line indicated the maximum possible performance of this agent in B in the 100:0 protocol. **E** Average choice ratios of individual flies from C showing significant learning in both 100:0 and 80:20 protocols (Wilcoxon signed-rank test: 100:0, p = 0.0039; 80:20, p = 0.0312) **F** Average choice ratios of individual flies from D. Flies showed a significant preference towards the rewarded odor in 100:0 but not 80:20 (100:0, p = 0.0391; 80:20, p = 0.1875). **G** Example fly data showing probability of accept odors as a function of odor experience number(blue), from flies performing the dynamic foraging protocol with DANs activated as reward.Fit using an MB-inspired regression model that incorporates either a covariance-based rule with reward expectation (gray), or a non-covariance rule (red). **H** Block-averaged choice ratios and reward ratios plotted against each other for flies with DAN activationused as reward (n = 26 flies, 3 blocks each). The best fit line for the DAN reward data is in blue. Best fit Gr64f sugar sensory reward behavior (Fig. 3C) is in red for comparison. **I** Block-averaged choice ratios and reward ratios (n = 50 simulations * 3, 80 trial blocks) from data simulated using a non-covariance-based rule are plotted against each other. The best fit line for the simulated data is in black, and for DAN reward (G) is in blue. **J** The instantaneous choice ratio of an example fly performing the dynamic foraging protocol plotted against trial number. Reward probabilities for trial blocks are indicated by the Y-arena icons above *(top)*. Individual odor choices are denoted by rasters, tall rasters - rewarded choices, short rasters - unrewarded choices. Curves show 10-trial averaged choice ratio and reward ratio, and horizontal lines the corresponding averages over the 80-trial blocks. **K** Change in percentage deviance explained, computed by subtracting the percentage deviance explained of the non-covariance-based model from a covariance-based rule, plotted for each fly (n = 26). On average, fly behavior was better predicted by the non-covariance-based model (Wilcoxon signed-rank test: p = 0.0164). Individual flies that were better fit by the covariance-based model have a positive value on this plot (gray region) while flies better fit by the non covariance-based model have a negative value (red region).

This observation that a non-covariance plasticity rule saturates at 75% preference in a 100:0 experiment and 50% in the 80:20 experiment is somewhat counter-intuitive. It arises because the non-covariance learning rule can only modify synapses in one direction - depression. In combination with the fact that a fly’s initial encounter with an odor is random (because we randomize right/left assignment of odors on each trial), we can account for the low preference of the non-covariance model. Consider the 100:0 case. Synapses related to the rewarded odor will have decreased to its minimum value, causing agents to always accept this odor. But agents only experience this odor 50% of the time. The other 50% of the time they experience the unrewarded odor. Agents accept and reject this unrewarded odor equally as they have never learnt anything in response to this. A first approximation would suggest therefore that this model would at max choosed the rewarded odor 75% of the time. A similar logic explains the 50% plateau performance in the 80:20 task. Given a sufficient number of reward pairings, responses to all odors will saturate at the most depressed state, and so the probability of accepting both cues becomes identical.

One potential concern with these experiments is that differences in optogenetic activation of the central DANs versus the peripheral Gr64f neurons could contribute to these behavioral differences. To address this, we tested whether similar levels of performance were achieved in a Pavlovian paradigm using these two different forms of optogenetic reward. Pavlovian learning was tested in a circular arena with LED intensity (2.3 mW/cm^2^) matched as closely as feasible to that used in the Y-arena (1.9 mW/cm^2^). Both PAM DAN and Gr64f sugar neuron activation support similar levels of learning performance in this assay (Supp. Fig. 9A-D) suggesting that optogenetic activation was sufficient to activate both types of neurons.

We next examined how bypassing reward expectation affects matching behavior. When tested with the same three-block matching design as earlier, but now providing a consistent reward signal via direct DAN stimulation, flies exhibited strongly diminished matching behavior (Fig. 5G,H). The slope of the choice-ratio, reward-ratio relationship was lower than that observed with Gr64f-driven reward, and approached the flat line predicted by simulations of behavior with a non-covariance based learning rule (Fig. 5I). The instantaneous choice ratio and reward ratio of an example fly (Fig. 5G) suggested that this flattening arises because choice ratios are never strongly biased to either odor. This is again explained by the uni-directional nature of the non-covariance rule. In agreement with this, changes in choice ratio at block transitions were much flatter with DAN reward than with Gr64f (Supp. Fig. 8C,D). To quantitatively evaluate whether providing reward with DAN activation changed the learning rule from covariance-based to a non-covariance rule, we fit our MB-inspired regression models (Fig. 4A) to fly data produced with DAN reward. We found that the non-covariance rule was the better fit (Fig. 5J,K). We find through these experiments that bypassing the computation of reward expectation changes fly choices from resembling behavior produced by a covariance-based learning rule, to behavior expected from a non-covariance rule. In particular, the results suggest this covariance-based rule is located in the fly MB and incorporates reward expectation but not sensory expectation.

All together, our results support the theory that covariance-based learning rules that incorporate reward expectation are necessary for operant matching. It suggests that reward expectation signal is calculated in the DANs of the fly MB and provides the first mapping of learning rules underlying operant matching onto plasticity mechanisms at a specific synapse.

## Discussion

The foraging strategies used by animals play a key role in their survival. Operant matching is one simple and ubiquitous behavioral strategy, utilized in dynamically-changing and probabilistic environments. Despite the ubiquity of this strategy and strong theoretical background, little is known about the underlying biological mechanisms. We leveraged the growing body of knowledge regarding learning in the fruit fly and the plethora of available anatomical tools to tackle this knowledge gap. We developed a foraging task that allowed us to monitor choices of individual fruit flies and showed, for the first time, that flies follow Herrnstein’s operant matching law. Combining experimental results with computational modeling, we found that this behavior requires synaptic plasticity and uses a rule that incorporates expectation of reward. Follow-up experiments manipulating neural circuitry found that reward expectation signals were incorporated via the rewarding PAM DANs. Our results provide the first mapping of the learning rule underlying operant matching onto the plasticity of specific synapses – the KC-MBON synapses in the MB.

### Does the ubiquity of operant matching imply a common mechanistic framework?

The operant matching strategy we observed in flies is wide-spread across the animal kingdom. When choosing between options that predict reward with different probabilities, mammals, birds and insects all follow Herrnstein’s matching law(11, 12, 14, 15, 17–19, 95, 96). This is clear at the global, trial-averaged level, where choice ratios are roughly equal to reward ratios, but is also true at the trial-by-trial level (Fig. 2A). In fact, we found that individual choices made by flies depended on choice and reward information received over multiple past trials (Fig 2G-H). This is in agreement with what has been observed in mice and monkeys(15, 19) and suggests that these animals all make use of similar kinds of information to guide their behavior. Flies also show an increase in speed of choice when rewarded, another common signature of learnt behavior in mice and monkeys(14, 15) (Supp. Fig. 1B,C).

It is unclear whether these behavioral similarities result from underlying mechanisms that are shared across species. At its surface, mechanistic similarities seem likely. For example, neural signals that subtract reward expectation from reward – a key component of the plasticity rules underlying matching shown here – can be found in the form of a reward prediction error in many different animals(6, 97–99). Nevertheless, such a signal on its own is not sufficient to produce matching; it needs to be incorporated into a covariance-based plasticity rule in a behaviorally relevant circuit. Note that mechanistic similarity would not require this specific use of reward prediction error, as covariance-based rules can also be implemented in a variety of other ways, as discussed at length in the Results. On the other hand, while learning values of options via synaptic plasticity is the traditional mechanistic framework thought to underlie such foraging decisions(21, 22), recent work has found signatures of graded neural responses proportional to value during inter-trial-intervals, suggesting a persistent-activity-based mechanism for foraging decisions that may not require synaptic plasticity(15, 100). Associated modeling efforts, including the work of Loewenstein and Seung, suggest that matching can arise from models that don’t incorporate synaptic learning(24, 101).

While both synaptic plasticity and non-plasticity mechanisms can explain the observed behaviors, each makes different testable assumptions about the underlying neural architecture(23) and the effect of changing environmental conditions on the behavior. For example, if one eliminated reward baiting in our experiment, a circuit using a covariance-based plasticity rule would still give rise to behavior that follows Herrnstein’s matching law. In this case, following such a law would lead the animal to always choose the option with higher reward probability. On the other hand, if matching behavior was produced using a different mechanism, the lack of reward-baiting might give rise to different strategies, such as the probability matching strategy commonly observed in mice under these conditions(8). Experiments to identify which mechanisms are used by different brains, and theoretical work to understand why, would therefore provide important insight into circuit function and the neural basis of operant matching.

### Plasticity in multiple MB compartments could explain deviations from matching

One complication to the framework of expectation-based learning rules and matching is that flies, like several other animals, don’t perfectly follow the matching law; rather they undermatch. Two hypotheses have been proposed to account for this deviation. The first proposes that animals that undermatch make use of a learning rule that deviates from a strictly covariance-based rule(26). One possibility for how this could occur is to have plasticity at multiple sites contributing to the overall learning, with different plasticity rules at each site. Indeed, the MB is divided into multiple compartments that contribute to behavior but exhibit important differences in learning(31, 34). It is possible that some compartments make use of reward expectation in a covariance-based learning rule, while others do not. Alternatively, undermatching can also result if reward expectations are estimated over long timescales(17), even if all compartments made use of a covariance-based rule. This idea suggests that in a dynamic environment where reward probabilities change quickly, the memory of past experiences acts as a bias that prevents the animal from correctly estimating the present cue-reward relationships. This is possible in the MB, as different compartments form and decay over different time scales(34). Whether either or both of these hypotheses explain undermatching in flies can be studied in future experiments by manipulating different compartments of the MB circuitry and analyzing the effect of such a manipulation on undermatching. Relatedly, it would be interesting to check animals adapt the timescales used to estimate reward expectations to the dynamics of the behavior task.

### An approach for inferring learning rules from behavior

Here we introduced a statistical method that uses logistic regression to infer learning rules from behavioral data. While we specifically applied our approach to infer learning rules for the fly mushroom body, the inference of learning rules is of importance to many areas of neuroscience(102, 103) this method could be similarly applied to model other learnt behaviors in the fly and other animals. In the current work, we considered learning rules that only depended on the current sensory stimulus (KC response) and reward (DAN response), but our methodology would also generalize to the inference of learning rules that incorporated a longer time-scale history of sensory input and reward. For example, the framework would be able to estimate rules that incorporated the weighted average of recent sensory experience.

However, it’s important to realize that the logistic regression formalization would break down entirely for learning rules that depend on the magnitudes of synaptic weights or postsynaptic activity. Such terms would induce different nonlinear dependencies between the choice sequence and learning rule parameters, preventing us from converting these choice and reward histories into regression inputs related to each component of the learning rule (see Methods). Our approach was appropriate here because the plasticity rule in the mushroom body was known to not involve these terms. However, many biological learning rules do depend on postsynaptic activity and current synaptic weights, and future work should explore more flexible methodologies from modern machine learning to develop generally applicable approaches.

### Circuit mechanisms for matching and reward expectation in *Drosophila*

We have shown that operant matching is mediated by synaptic plasticity in the fly mushroom body that requires the calculation of a reward expectation. However the mechanisms underlying this calculation remain unclear.

The proposed mechanism underlying the calculation of reward prediction error (RPE) in mammals provides a hint at one option(97, 104). Here dopaminergic neurons implicitly represent expectation by calculating the difference between the received reward and the reward expectation. This has been found to involve the summation of positive ‘reward’ inputs and negative GABA-ergic ‘expected reward’ inputs to the dopaminergic neurons(105). MB DANs could represent reward expectation in a similar way. In fact, the recently released hemi-brain connectome(44) has found many direct and indirect feedback connections from MBONs to DANs that theoretical work has shown could support such a computation(65, 94). In the MB circuit, MBON activity is related to the expectation of reward associated with a given odor(31, 56). An inhibitory feedback loop, via GABA-ergic interneuron(s) for example, could potentially carry reward expectation related information from MBONs to DANs. The negative expected reward signal from this interneuron could be combined in the DANs with a positive reward signal from sensory neurons, allowing DAN activity to represent the type of reward expectation signal needed by a covariance-based rule.

It is important to note that such a mechanism would have a major difference from mammalian RPEs. Since MBON activity is linked to the presence of odor, the reward expectation signal would vary across stimuli and only be present when the stimulus was too. Thus, this signal would not have the temporal features of mammalian RPEs. This difference in temporal structure of the reward expectation signal could explain the mixed observations from past studies aimed at identifying reward expectation in flies. For instance, a study that used temporally distinct cues and reinforcements suggested that DANs do not incorporate reward expectation(68), while studies that used temporally overlapping cues and reinforcements did find signatures of reward expectation(69, 72), albeit with different temporal properties than the typical mammalian RPE.

It’s also possible that reward expectations are incorporated into mushroom body plasticity by adjusting the levels of reward and punishment needed to achieve a given dopamine signal. In this scheme reward-related dopamine neurons could represent how much a reward exceeds expectations, and punishment-related dopamine neurons could respond when expectations are not met. This is reminiscent of the idea from Felsenberg and colleagues that interactions between reward and punishment-related compartments in the MB can guide bi-directional learning(36, 66, 67, 71). However, here we extend the idea by proposing that reward would not only modify KC-MBON synapses, but also modulate the baseline dopamine release or firing threshold of reward-related dopaminergic neurons. Similarly, upon missing an expected reward, learning would do the same for MBONs and DANs in punishment-related compartments. The resulting behavior would depend on balance between the activity of both reward and punishment compartments, and if the reward and punishment baselines were updated correctly, such a mechanism could produce a covariance-based rule and support operant matching. This mechanism would also tie into the notion that phasic dopamine release (i.e. the difference of dopamine from its baseline level) mediates the RPE signal in mammals(106).

Future experiments can distinguish between these mechanistic hypotheses. For instance, neural recordings can probe how DAN activity changes over the course of the task, and connectomics can identify other neurons in the system that may be important for the computing reward expectation. These types of experiments are easily doable in the *Drosophila melanogaster* model. Paired with further modeling efforts and the foraging framework we developed, the fly MB promises to be a system in which we can understand decision making at a level of detail that is presently unparalleled in the field of systems neuroscience.

## Materials & Methods

### Fly strains and rearing

*Drosophila melanogaster* used for behavior experiments were raised on standard cornmeal food supplemented with 0.2 mM all-trans-retinal at 25°C [for Gr64f-Gal4 lines - see table] or 21°C [for other lines - see following table] with 60% relative humidity and kept in dark throughout. The details of all flies used for experiments in this manuscript can be found in the table below:

| Genotype | Expression target/reporter description | Bloomington stock number /Reference (if applicable) |
| --- | --- | --- |
| w; Gr64f-Gal4/CyO;<br>Gr64f-Gal4/TM3 X<br>20XUAS-CsChrimson-mVenus attp18 | Optogenetic activation of Gr64f expressing sugar sensory neurons in behavior experiments | (Dahanukar et. al. 2007; Haberkern et. al. 2019) (84, 107)<br>X<br>BDSC:55134(Klapoetke et. al. 2014) (41) |
| w; +; 58E02-Gal4<br>X<br>20XUAS-CsChrimson-mVenus attp18 | Optogenetic activation of PAM cluster DAN in behavior experiments | BDSC:41347(Jenett et. al. 2012) (40)<br>X<br>BDSC:55134(Klapoetke et. al. 2014) (41) |
| w; 10XUAS-opCas9wt in ZH51C; DopR1-4gRNA JK65C/CyO::TM6B<br>X<br>w; +; 13F02-Gal4 attP2, Gr64LexAp65 JK73A, 13XLexAop-IVS-Syn 21-Chrimson88::tdT/TM3 | Optogenetic activation of Gr64f expressing sugar sensory neurons and CRISPR mediated knockout of DopR1 receptors in KCs for behavior experiments | New Stocks - see Cloning sub-section below |
| w; 10XUAS-opCas9wt in ZH51C; DopR1-4gRNA JK65C/CyO::TM6B<br>X<br>w; +; Gr64LexAp65 JK73A, 13XLexAop-IVS-Syn 21-Chrimson88::tdT/TM3 | Optogenetic activation of Gr64f expressing sugar sensory neurons without expression of cas9 protein needed for knockout of DopR1 in behavior experiments | New Stocks - see Cloning sub-section below |

Cross progeny (2-5 days old) were sorted on a cold plate at around 4°C and females of the appropriate genotype were transferred to starvation vials. Starvation vials contained nutrient-free 1% agarose to prevent desiccation. Flies were starved between 28 - 42 hrs before being aspirated into the Y-arena for experiments.

#### Cloning

The Gr64f promoter was PCR amplified using Q5 High-Fidelity 2× Master Mix (New England Biolabs) from the Gr64f-GAL4 (107) and cloned into the FseI/ EcoRI digested backbone of pBPLexAp65 (39) using NEBuilder HiFi DNA Assembly (New England Biolabs). Primer sequences were:

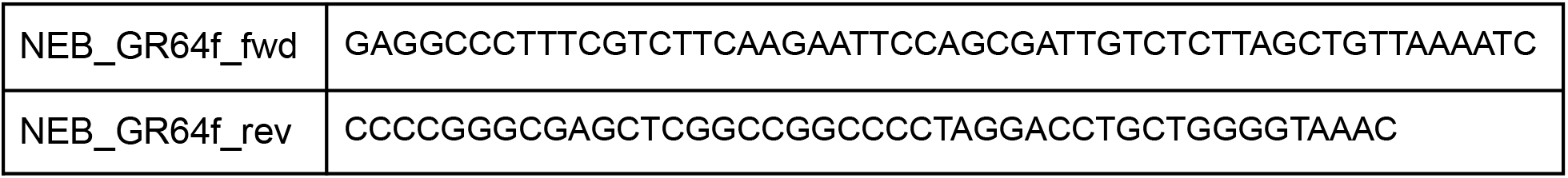

Four gRNA for the gene Dop1R1 were designed using https://flycrispr.org/target-finder/ (89). The gRNA were then cloned into pCFD5_5 following the protocol published in Port and Bullock, 2016(108).

Dop1R1 gRNA target sites (5’-3’)

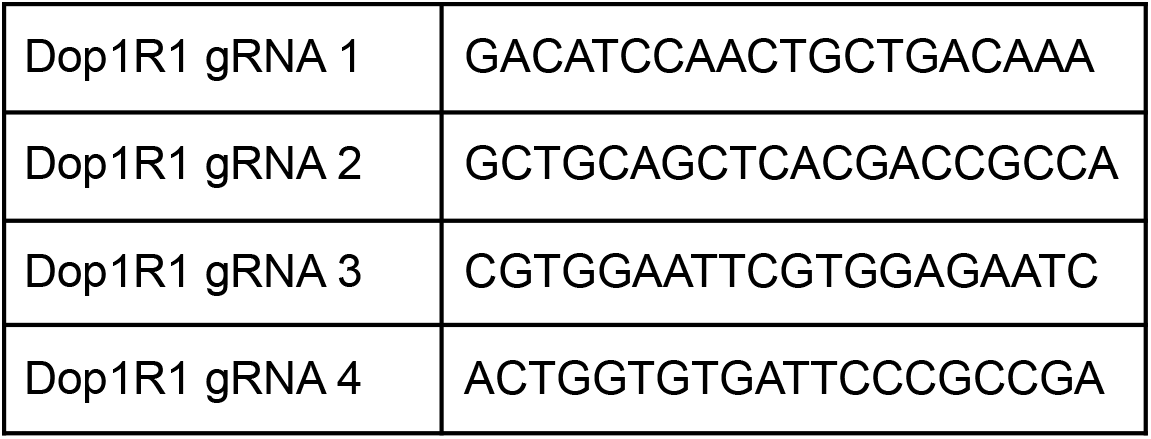

Primer sequences were:

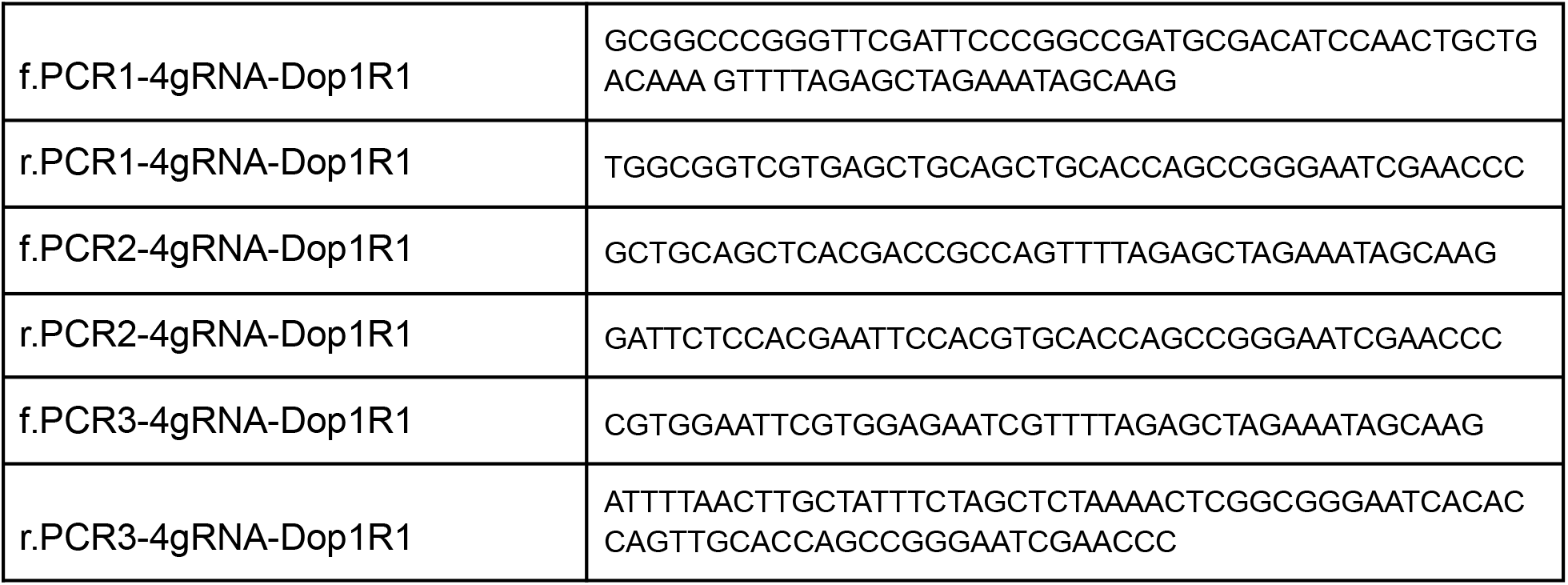

Transgenic injections were performed by Genetivision using fC31 integrase mediated integration into attP dock sites. Gr64f-LexAp65 was integrated into P{CaryP}JK73A and Dop1R1 was integrated into P{CaryP}JK65C.

### Y-arena

Single fly behavior experiments were performed in a novel olfactory Y-arena designed in collaboration with the Janelia Experimental Technology team (jET).

#### Apparatus design

A detailed description of the apparatus is provided in Supplementary Information 1. The Y chamber consists of two layers of white opaque plastic. The bottom is a single continuous circular layer and serves as the floor of the Y that flies navigate. The top is a circular layer with a Y shaped hole in the middle. The length of each arm from center to tip is 5 cm and the width of each arm is 1 cm. These two layers are placed underneath an annulus of black aluminum. A transparent glass disk is located in the center of this annulus and acts as the ceiling of the Y - allowing for video recording of experiments. This transparent disk is rotatable and contains a small hole used to load flies. The black annulus houses three clamps that hold the circular disk in place. All three layers are held together and made airtight with the help of 12 screws that connect the layers.

The Y chamber is mounted above an LED board that provides infrared illumination to monitor the fly’s movements, and red (617 nm) light for optogenetic activation. The LED board consists of a square array of red (617 nm peak emission, Red-Orange LUXEON Rebel LED, 122 lm at 700mA, 1.9mW/cm^2^) and infrared (IR) LEDs that shine through an acrylic diffuser to illuminate flies. Experiments were recorded from above the Y using a single USB3 camera (Flea3, model: FL3-U3-13E4M-C: 1.3 MP, 60 FPS, e2v EV76C560, Mono; Teledyne FLIR, with longpass filter of 800 nm).

Each arm of the Y has a corresponding odor delivery system, capable of delivering up to 5 odors. For our experiments, olfactometers injected air/odor streams into each arm at a flow rate of 100 ml/min. A crisp boundary between odors and air is formed at the center of the Y (Supp. Fig. 1A). Odors and concentrations used for each experiment are detailed in the behavioral task structure and design section of the methods. The center of the Y contains an exhaust port connected to a vacuum, which was set at 300 ml/min using a flow meter (Dwyer, Series VF Visi-Float® acrylic flowmeter) - matching total input flow in our experiments.

#### Fly tracking and operation

We wrote custom MATLAB code (MATLAB 2018b, Mathworks) to control the Y-arena and run experiments. The data collected by the USB3 camera was loaded into MATLAB in real time and the fly’s location was identified using the MATLAB image processing toolbox as follows. A background image was calculated just before beginning the experiment by averaging multiple frames as the fly moved around in the Y. This background was subtracted from the frame being processed and the resulting image was thresholded, leaving the fly as a white shape on a black background. The location of the centroid of the fly was estimated using the MATLAB’s *bwconncomp* and *regionprops* functions and then assigned to 1 of 6 regions in the Y. If the fly was located in one of the reward zones, a trial was deemed complete and rewards were provided by switching on the red LEDs as defined by the reward contingencies of the task. The arena was then reset with air being pumped into the chosen arm and odors randomly reassigned to the two other arms (Fig. 1B). The location of the fly along with other information, such as reward presence and odor-arm assignments, were saved as a .mat file for further analysis. All analysis in Figures 1, 2 and 3 were based on this saved information.

### Circular olfactory arena

Group learning experiments in Supp. Fig. 9 were performed in a previously described circular olfactory arena(31).

### Behavioral experiments

#### Odorant information

For all experiments in the paper, two or three of the following odorants were used to form cue-reward relationships:

1. 3-Octanol (OCT) [Sigma-Aldrich 218405]. *Y-arena:* diluted in paraffin oil [Sigma-Aldrich 18512] at a 1:500 concentration and then air-diluted to a fourth of this concentration. *Circular olfactory arena:* diluted in paraffin oil at a 1:1000 concentration in the circular arena.
2. 4-Methyl-cyclo-hexanol (MCH) [Sigma-Aldrich 153095]. *Y-arena:* diluted in paraffin oil [Sigma-Aldrich 18512] at a 1:500 concentration and then air-diluted to a fourth of this concentration. *Circular olfactory arena:* diluted in paraffin oil at a 1:750 concentration in the circular arena.
3. Pentyl Acetate (PA) [Sigma-Aldrich 109584]. *Y-arena:* diluted in paraffin oil [Sigma-Aldrich 18512] at a 1:5000 concentration and then air-diluted to a fourth of this concentration

#### Y-arena behavioral task structure and design

In the 100:0 protocol flies were inserted randomly into one of the three arms. This arm was injected with a clean airstream and OCT and MCH were randomly assigned to the other two arms. For a given fly, one of OCT or MCH was paired with reward 100% of the time. Once a fly reached the choice zone of either the odor arm a choice was considered to have been made. If the rewarded odor was chosen, the fly was rewarded with a 500ms flash of red LED (617 nm, 1.9mW/cm^2^) to activate the appropriate reward-related neurons. The arena was then reset with the arm chosen by the fly injected with clean air and OCT and MCH randomly assigned to the other two arms. This was repeated for many trials. In the version of this task seen in Fig. 1C, flies were allowed to choose between OCT and MCH for 30 minutes when neither option was rewarding and then for 30 minutes when one of the options was consistently rewarding. In the version of this task used for analysis in Fig. 1B-D, and Fig. 5A-F, flies made 60 naive choices where neither option was rewarding and 60 training choices where one option was consistently rewarded. In the version used for analysis in Fig. 1E flies made 60 rewarded choices where one option was consistently rewarded, and in Supp.1B,D-F flies made 120 rewarded choices where one option was consistently rewarded.

In a probabilistic version of this task used in Fig. 1F, Supp. Fig. 1C, and Fig. 5A-F, rather than one of the options being consistently rewarded and the other not, both options were rewarded probabilistically. The sum of reward probabilities for both options for this version of the task was either 1 or 0.5. Whenever probabilistic rewards were included in our tasks, reward baiting was also incorporated as follows. If an unchosen action would have been rewarded, the reward was delivered on the next choice of that alternative with 100% certainty. This means that the likelihood an odor cue holds a reward increases over time if it is unchosen for many trials. Importantly, the fly performing the task is never informed as to whether the unchosen option would have been rewarding on any given trial.

In Fig. 1G and Supp. Fig. 1G, we analyzed a version of this task where flies learnt multiple simultaneous probabilistic cue-reward pairings. Here every fly experienced all three odors. For a given fly OCT and MCH arbitrarily paired with reward 80% and 40% (or 40% and 80%) of the time, respectively. EL was unpaired. Any given trial of these experiments consisted of a choice between OCT and EL or MCH and EL. These two options were interleaved together. Flies experienced 40 naive trials of each combination where no rewards were provided. This was followed by 40 rewarded trials.

The dynamic foraging task in Fig. 2 was adapted from monkey and mouse versions (13–15, 18) and incorporated features such as baiting and probabilistic reward described earlier. OCT and MCH were the two odors used in this task. An example protocol of this task with details about the three block structure and number of trials per block can be seen in Fig. 2A. The relative reward ratios between the two odors for a given block were drawn from the following set [1:1, 1:2, 1:4, 1:8]. The sum of reward probabilities for both options for this version of the task was either 1 or 0.5.

#### Circular olfactory arena behavioral task structure and design

A schematic of the task performed in the circular arena is shown in Supp. Fig. 9A. OCT and MCH were used as odors for these experiments. Odors were presented sequentially and separated in time for one minute each, with one of the odors paired with reward. To mimic the relationship between odor time and reward time experienced by the fly in the Y-arena, 1 sec of reward (red light, 617nm, 2.3mW/cm^2^), was provided for every 3 seconds of odor experience. Flies were finally tested by dividing the circular arena into four quadrants with two opposite quadrants receiving one odor and the other two quadrants receiving the other.

### Quantitative analysis and behavioral modeling

All analysis and modeling was performed using MATLAB 2020b (Mathworks).

#### Analysis of fly location in the circular arena

Videos of a fly’s movements in the Y-arena were read into MATLAB frame by frame and the location of the fly’s centroid was identified using the MATLAB image processing toolbox. Once identified, the number of flies in each quadrant was used to calculate the preference index (PI) metric. PI is defined as the difference between the number of flies in each pair of odor-matched quadrants divided by the total number of flies.

#### Analysis of fly movement and choices in the Y-arena

The *(x,y)* coordinates of the fly were analyzed to calculate: i) the distance of the fly from the center of the Y; ii) when the fly entered and exited a given odor arm; and iii) the time taken per trial to enter into the reward zone at the end of an odorized arm. These quantities were then used to produce the plots in Fig. 1, 2, 5 and Supp. Fig. 1, 2.

Distance from center was calculated by projecting the (*x,y*) location of the fly (*p*_*t*_) onto a skeleton of the Y. Here the subscript *t* denotes the time point at which the (*x,y*) location was observed. The skeleton consisted of three lines running down the middle of each arm to the center of the Y (*υ*_0_). Based on which arm the fly was located in, its (*x,y*) position was projected onto the appropriate (*i*^*th*^) skeleton line using the following equations for projecting a point onto a line:

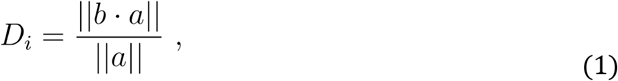

where *b* = *p*_*t*_ − *υ*_0_, *a* = *υ*_*i*_ − *υ*_0_, and *υ*_*i*_ is the (x,y) coordinates of the end of the *i*^*th*^ arm. The entries/exits of a fly into/from a particular odorant or air were estimated by tracking the region that the fly was located in at every time point and comparing it to the known odor-arm identity map (stored in the experiment .mat file). A turn (reversal) was considered to have been made whenever a fly entered an odor and then exited this odor without reaching the reward zone. An approach was considered to have been made whenever a fly entered an odor arm and then traveled all the way into the reward zone of that same arm without ever exiting it.

To calculate the time taken per trial, we made use of the timestamp vector that we saved along with the (*x,y*) vector. Time taken from the entire trial was calculated by subtracting the timestamp for the frame that the previous trial was completed by the timestamp of the frame when the current trial was completed. Time taken from first exit of the air arm was calculated by subtracting the timestamp of the frame that the fly first exited the air arm after a trial began by the timestamp of the frame when the current trial was completed.

Choices themselves were determined by identifying the arm in which the fly crossed into the reward zone and mapping that arm to its assigned odor on that trial.

#### Logistic regression to estimate influence of past rewards and choices on behavior

To estimate the role of choice and reward histories in determining fly choices in the dynamic foraging task, we fit the following logistic regression for each fly as

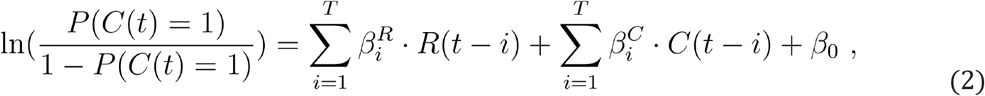

where *t* is the present trial and *i* is the variable used to iterate over the past *T* trials. *C*(*t*) = 1 if the chosen odor was OCT and -1 if the chosen odor was MCH. *R*(*t*) = 1 if chosen OCT option produced reward, -1 if chosen MCH option produced reward, and 0 otherwise. *β*_0_ represents the weight assigned to the bias term, 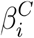 represents the weight assigned to the *i*^*th*^ past choice and 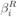 represents the weight assigned to the *i*^*th*^ past reward. We chose to look at the past *T* = 15 trials to align with previous studies (15, 19). The regression coefficients generated were 10-fold cross-validated, and the regression model included an elastic net regularization (MATLAB function - lassoglm). The weight of lasso versus ridge optimization was set to 0.1 as this value provided best behavioral fits. These fly-specific regression coefficients could be combined with the flies reward and choice histories to predict trial choice probability and estimate the log-likelihood (*l*) and percent deviance explained (*PD*):

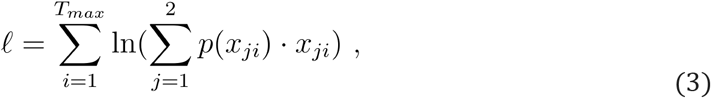

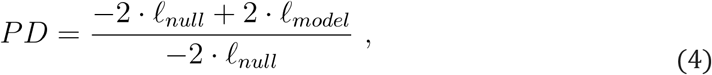

where *T*_*max*_ is the total number of trials in the data being fit, *i* indexes trials, *j* indexes possible options, *p*(*x*_*ij*_) is the probability with which the model predicts that choice *j* occurs on trial *i*, and *x*_*ij*_ is the choice that actually took place on trial.

#### Leaky-integrator model

We also developed a leaky-integrator model to predict behavior in the dynamic foraging task inspired by earlier work (18). This model determines choices on a given trial by comparing values assigned to each option the agent has to choose between based on choice and reward history.

The values (*Q*)were calculated for a given trial *t* using the following equations. If OCT is chosen by the model, values are updated according to

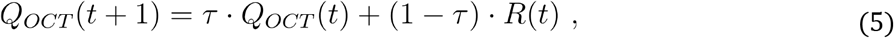

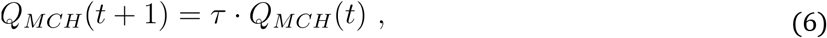

where *τ* is a constant related to the learning rate. Similarly, if MCH is chosen by the model, values are updated according to

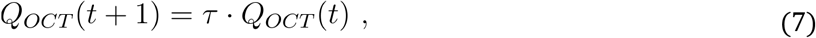

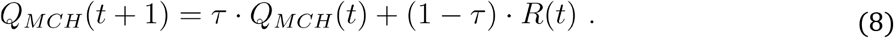

These values are then compared and passed through a sigmoidal nonlinearity to determine the probability of each choice,

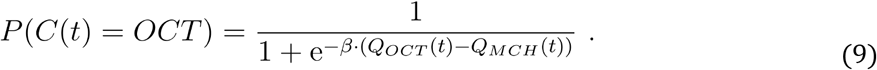

The probability of choosing MCH was one minus that of OCT. The probability generated by this operation is compared with a value drawn from a uniform distribution over the [0,1] interval to determine whether the resulting choice is OCT or MCH. These predicted choices could be compared to fly behavior to compute the model’s fraction deviance explained. The parameters *β* and *τ* are fit for each fly so as to maximize the percentage deviance explained (values of these parameters can be seen in Fig 2G and Supp. Fig. 3A,B).

#### Win - stay, Lose - switch model

A third model to predict behavior incorporated information only about the most recent choice made by the fly unlike the logistic regression and leaky-integrator alternatives. In this “stay-switch” model the agent chooses randomly on the first trial. If the chosen option produces a reward the agent picks the option again on the next trial (stays). If it doesn’t produce a reward the agent picks the other option on the next trial (switches). This procedure repeats to generate a sequence of choices. The accuracy of this model was calculated by observing correctly predicted switches and stays as well as incorrectly predicted switches as stays (Supp. Fig. 3E).

### Neural circuit model of dynamic foraging

We designed a neural circuit model, inspired by work from Loewenstein and Seung (24), that was used to simulate behavior in a dynamic foraging task. Two versions of this model were used.

### Replicating Loewenstein and Seung’s Model

The first version aimed to directly replicate the model used by Loewenstein and Seung (Supp. Fig. 4A). It generated behavior on a trial-by-trial basis in the dynamic foraging task. The number of trials to simulate were input by the user prior to simulation (typically 60, 240 or 2000 trials). The model consisted of two sensory neurons (*S*_1_ and *S*_2_) whose activity was drawn at the beginning of each trial from a normal distribution with mean 1 and standard deviation 0.1. These neurons synapse with weights (*W*_1_ and *W*_2_) onto two motor neurons (*M*_1_ and *M*_2_), whose activity was given by the weighted sum of sensory neuron activity. The activity of *M*_1_ and *M*_2_ were compared and the choice was driven by whichever neuron had the larger activity.

Once the choice was made, rewards were provided as determined by the reward contingencies of the task. These were input by the user prior to running the simulation. The weights between *S* and *M* were updated after each choice and followed the following rules

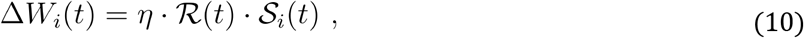

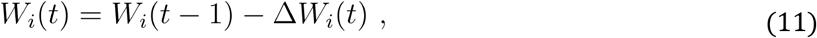

where ℛ = *R* or ℛ = *R* − *E*(*R*), 𝒮_*i*_ = *S*_*i*_ or 𝒮_*i*_ = *S*_*i*_ − *E*(*S*_*i*_) based on the learning rule, and *i* iterates over odors. Note that *E*(*R*) and *E*(*S*_*i*_) depended on time and were calculated in one of two ways: i) by calculating the mean over the last 10 trials, ii) by filtering the entire history with an exponential filter with exponential timescale(*τ*)of 3.5 trials. The various covariance and non-covariance rules were achieved by selecting the appropriate combinations of ℛ and 𝒮_*i*_.

### Task and Mushroom-body Inspired Version

The second version incorporated modifications to the model that made it more appropriate for the task we designed for fruit flies (Fig.3B). This model consisted of two sensory inputs that represented activity of populations of Kenyon cells (KCs). However, this version of the model looped through odor experiences, rather than looping through trials determined by two-alternative forced choices. Therefore the activity of the sensory neurons was drawn differently. Rather than both values being drawn from normal distribution with mean 1 and standard deviation *σ* = 0.1, this was only true for the odor that was deemed to have been “experienced” by the model on a given odor experience. The activity of the other neuron was drawn from a normal distribution with mean *α* = 0. 1 (Fig. 3,4; Supp. Fig. 5,6) and standard deviation *σ* = 0. 1 Here *α*, represents the similarity, or overlap, between the two inputs. This was included because the KC representations of the two different odors used in our task are thought to have some amount of overlap (92). However, we found that modulating this term did not affect the resulting matching behavior (Supp. Fig 6D-G) and so for Fig. 5, we chose *α*= 0. We also explored incorporating noise covariance between the two sensory inputs (with correlation coefficient *c* = 0.1), but this correlation was empirically unimportant and we usually set *c* = 0.

Another difference is that an odor experience could lead to either an approach (choice) or a turn away. The behavior chosen by the model on any given odor experience depended on the response of the single output neuron incorporated into this model. The activity of this output neuron (*M*) was the weighted sum of the two inputs. This was then passed through a sigmoidal nonlinearity

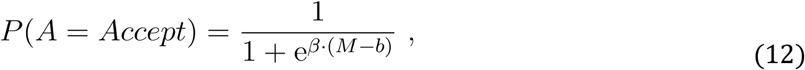

where *β* = 4, *b* = 1(this value was chosen to encourage exploration at the beginning of learning) and *A* is the action produced by the model. When *A* = 0 the odor is accepted and when *A* = 1 the odor is rejected. A random number from the interval [0,1] was drawn and compared to Y to determine whether an approach/choice or turn was made. If a turn was made, no reward was provided, and the weights remained unchanged. The model then experienced a new odor and the process repeated. If a choice was made then a reward was provided based on the choice contingencies and weights were updated according to the rules in eqs. 10 and 11.

### Logistic regression model for estimating learning rules

To determine the learning rules that best predict fly behavior, we designed a logistic regression model that made use of the known relationship between MBON activity and behavior. This model predicted behavior between input and weights that give rise to MBON activity following the relationships

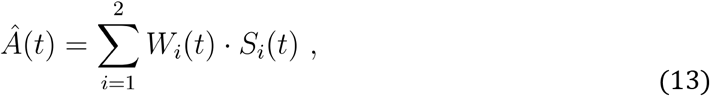

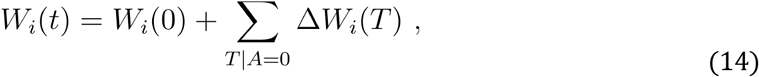

where *Â*(*t*) is the predicted action on odor experience *t, T*|*A* = 0 indicates all past odor experiences where the fly chose to accept the odor, and *W*_*i*_(*t*) represents the synaptic weights associated with neurons representing odor *i* at time *t*. Now the change in synaptic weights Δ*W*_*i*_(*t*) depends on the learning rule that is used by the circuit. It was here that we wanted to have the regression model identify the rule that provided the best fit to actual data. To do this we allowed the model to use a learning rule with 4 different terms whose coefficients could be modified,

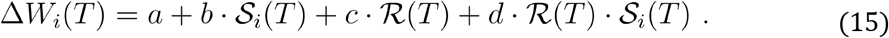

Here, *a, b, c*, and *d* are the coefficients assigned to each component of the learning rule. The regression model takes the sensory stimuli and synaptic weights at a given time as inputs to predict the output action. However, when fitting this model to behavior we have only sensory stimulus and reward information readily available. We therefore used eq. 14 and 15 to convert synaptic weights and sensory stimuli to inputs that consisted of sensory stimuli and rewards and a constant input that serves as a bias term. The resulting inputs could be represented as

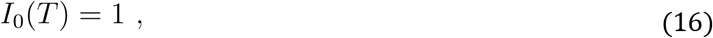

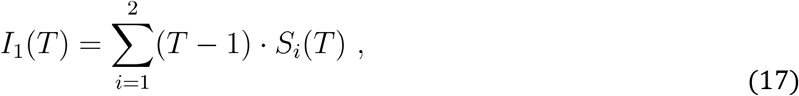

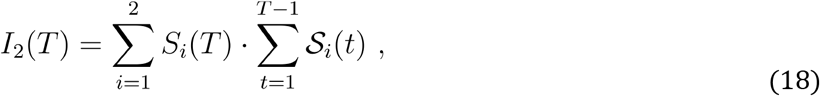

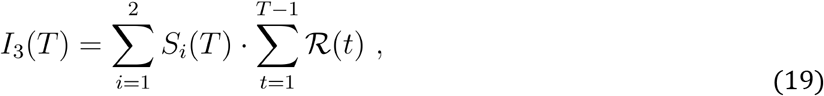

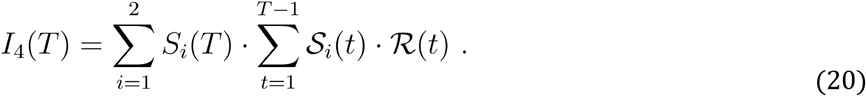

The coefficients assigned to each of the five inputs could then be used to identify the learning rule that the model predicted as the best estimate for producing the behavior that was tested.

## Acknowledgements

This work was supported by the Howard Hughes Medical Institute. We thank Igor Negrashov, Tobias Goulet, Peter Polidoro, Steven Sawtelle, Jon Arnold and others at jET, Janelia Research Campus for helping design and fabricate the Y-arena. We thank Todd Laverty and others in the Janelia Drosophila resources team for providing support with fly lines and media. We also thank all the members of the Turner, Fitzgerald and Aso groups for support and insightful discussions as well as Brad Hulse, Eyal Gruntman, Sandro Romani, Mehrab Modi, Yichun Shuai, Laura Grima, Luke Coddington and Yoshi Aso for feedback on the manuscript. AER would like to thank The Neuroscience Graduate Training Program at The Solomon H. Snyder Department of Neuroscience, Johns Hopkins University and thesis committee members Vivek Jayaraman, Ann Hermundstad, Jeremiah Cohen, Christopher Potter, Yoshi Aso and Erik Snapp for their guidance and support.

## SUPPLEMENTARY MATERIAL FOR

**Supplementary Figure 1:**
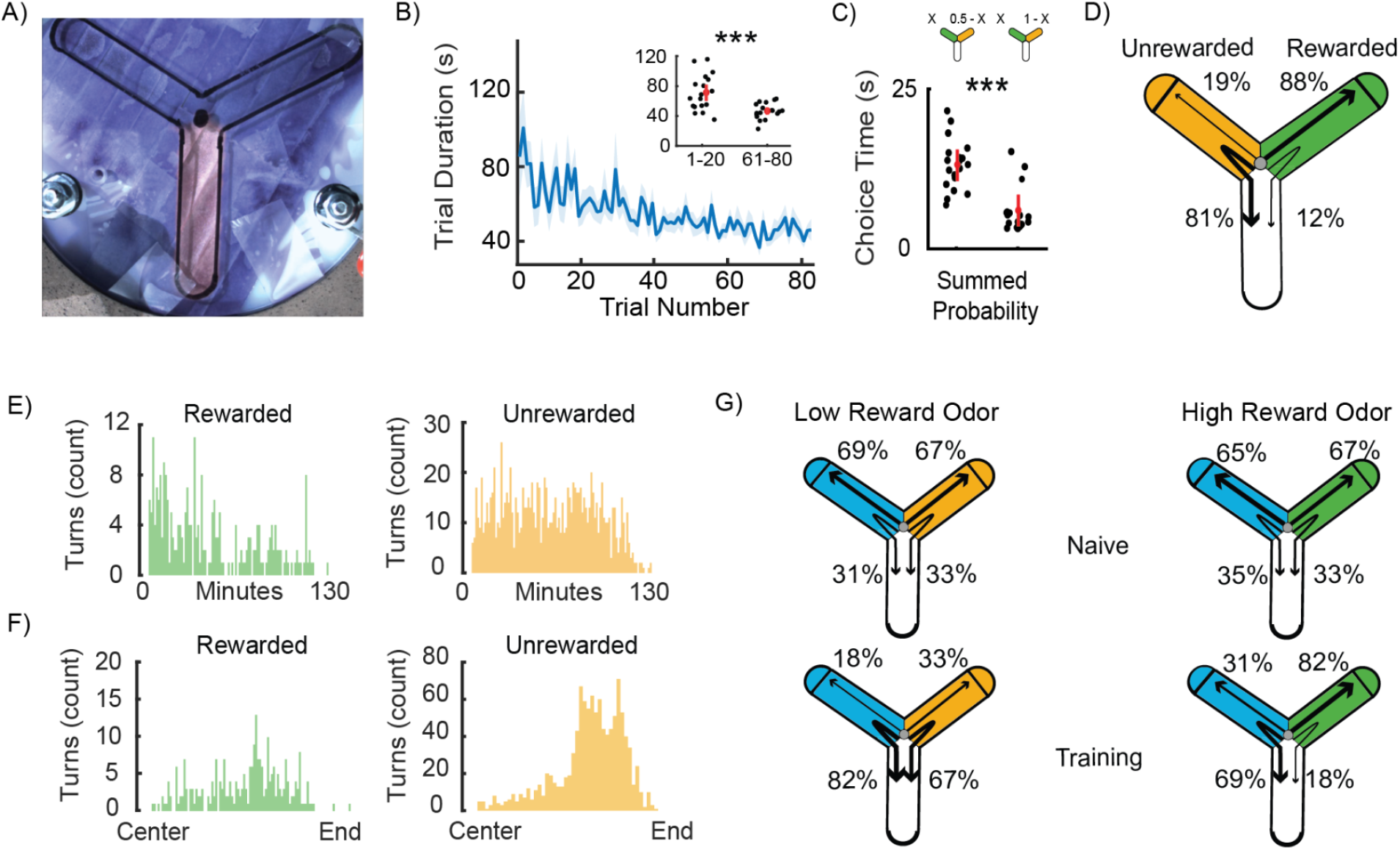
Further quantification of learning of multiple probabilistic cue-reward associations in the novel Y-arena. **A** A measurement of odor boundaries in the Y-arena. Moist blue litmus paper was placed in the Y-arena while the arm at the bottom was filled with carbon dioxide. This caused the color of the litmus paper to change, providing an estimate of how odor boundaries are formed at the center of the Y. **B** The time taken to make a choice decreases once reward is made available to the fly (mean +/- SEM). *Inset:* Average trial time for the first 20 trials is longer than the average trial for the last 20 trials (Wilcoxon signed-rank test: p = 6.292*10^−4^, n=18 flies). **C** The average choice time across 80 trials, measured from first exit of the air arm till entry into the reward zone, for two different summed probabilities of receiving reward (0.5 or 1). Average choice time decreases as the summed probability increases (Mann-Whitney rank-sum test: p = 4.7653*10^−4^, n = 18 for summed probability = 1; n = 20 for summed probability = 0.5). **D** Percentage of odor arm entries that lead to accept or reject in a 100:0 protocol. Flies show increased rejection of the unrewarded odor (Mann-Whitney rank-sum test: p = 3.2278 * 10^−7^, n = 18), and decreased rejection of the rewarded odor (Mann-Whitney rank-sum test: p= 3.2166 * 10^−7^, n = 18). **E** Histogram showing the number of rejects over time for rewarded and unrewarded odor choices. Rejects decrease over time for the rewarded odor (Wilcoxon signed rank test: p =0.0171, n = 18, comparing bins 15-65 with 66-115 minutes - to exclude large time values when many flies had already finished the task), but not the unrewarded one (Wilcoxon signed rank test: p = 0.1839, n = 18). **F** Histogram showing the number of reversals as a function of distance along the odorized arm. The demarcation of “End” on the x-axis represents entry into the reward zone. **G** The average percentages of accepting and rejecting each odor - high-rewarding (green), low-rewarding (orange) and unrewarded (blue) are graphically represented in a schematic of the Y-arena (n=10 flies). Flies increasingly accepted the high-rewarding odor(Mann-Whitney rank-sum test: p= 1.8267*10^−4^, n = 10), and displayed an increased probability of rejecting both low-rewarding and unrewarded odors, as compared to naive trials (Supp. Fig. 1G left; Mann-Whitney rank-sum test: unrewarded odor: p = 1.8165*10^−4^, n = 10; rewarded odor: p = 7.6854*10^−4^, n = 10).

**Supplementary Figure 2:**
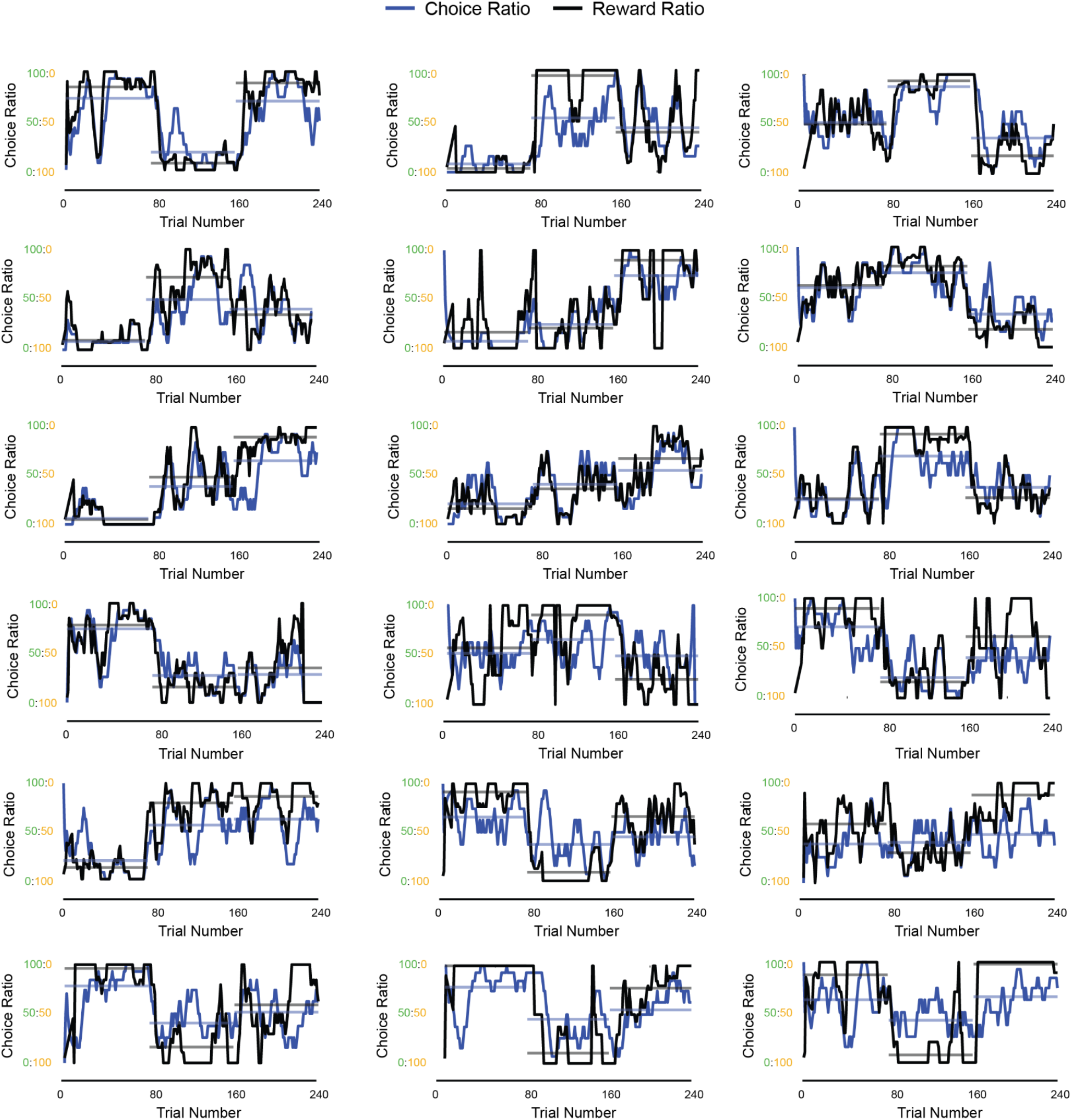
All example instantaneous choice ratio and reward ratio plots. Matching of instantaneous choice ratio (blue) and reward ratio (black) in all example flies. Curves show 10-trial averaged choice ratio and reward ratio, and horizontal lines the corresponding averages over the 80-trial blocks.

**Supplementary Figure 3:**
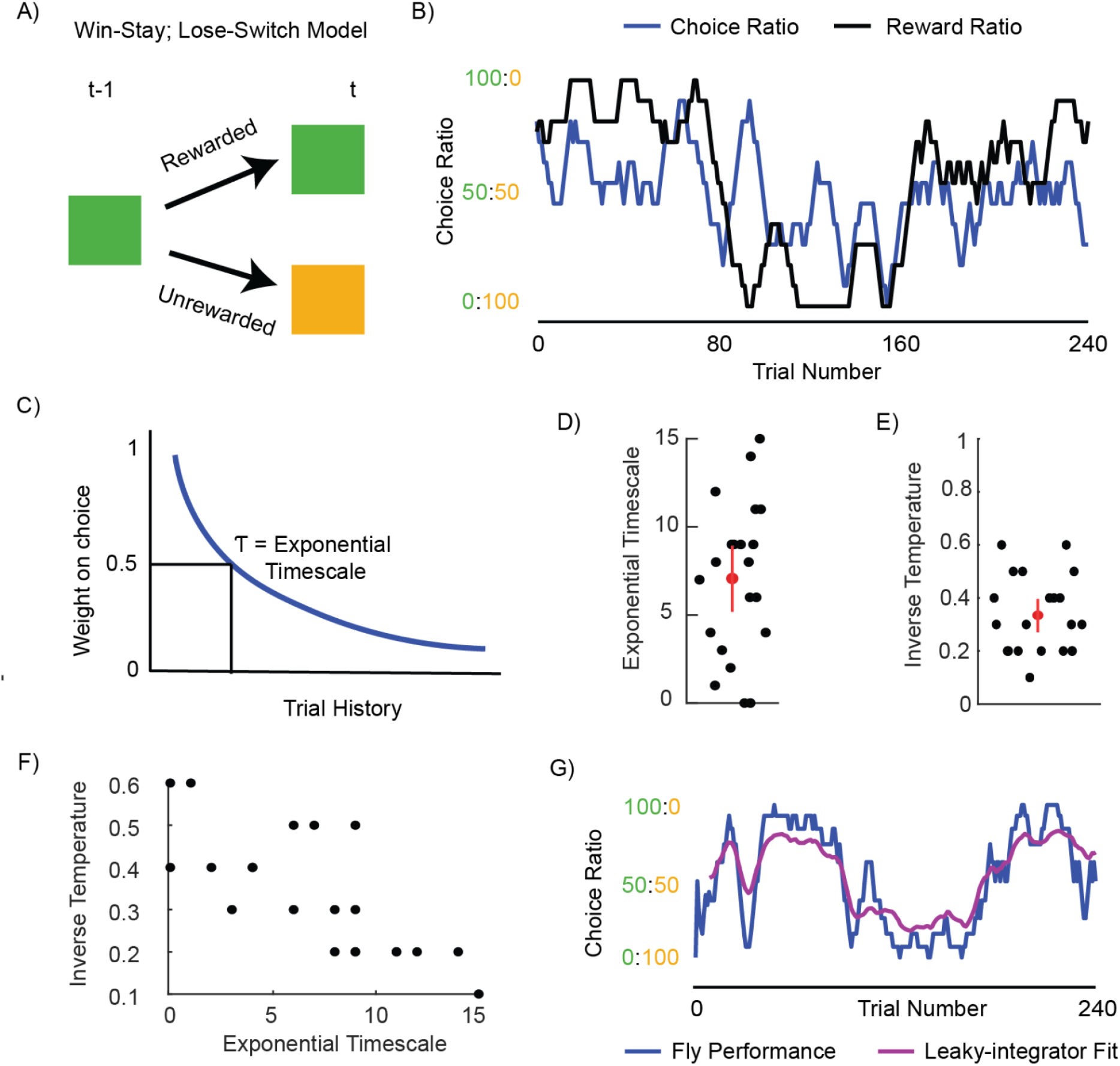
Analysis of the “win-stay lose-switch” and “leaky-integrator” models. **A** Schematic of the “win-stay; lose-switch” model **B** Example choice data generated by the “win-stay; lose-switch” model showing instantaneous choice ratio (blue) and reward ratio (black) **C** Schematic representing how the past trial history is weighted to calculate value in the “leaky-integrator” model. **D** Estimated exponential timescales (τ) for each fly shown in Supp Fig. 2 **E** Estimated inverse temperatures for each fly shown in Supp Fig. 2 **F** Relationship between exponential timescale and inverse temperature. **G** Leaky-integrator model fit (purple) on behavior (blue) from the example fly in Fig. 2A, plotted from the 15th trial onwards to avoid edge effects.

**Supplementary Figure 4:**
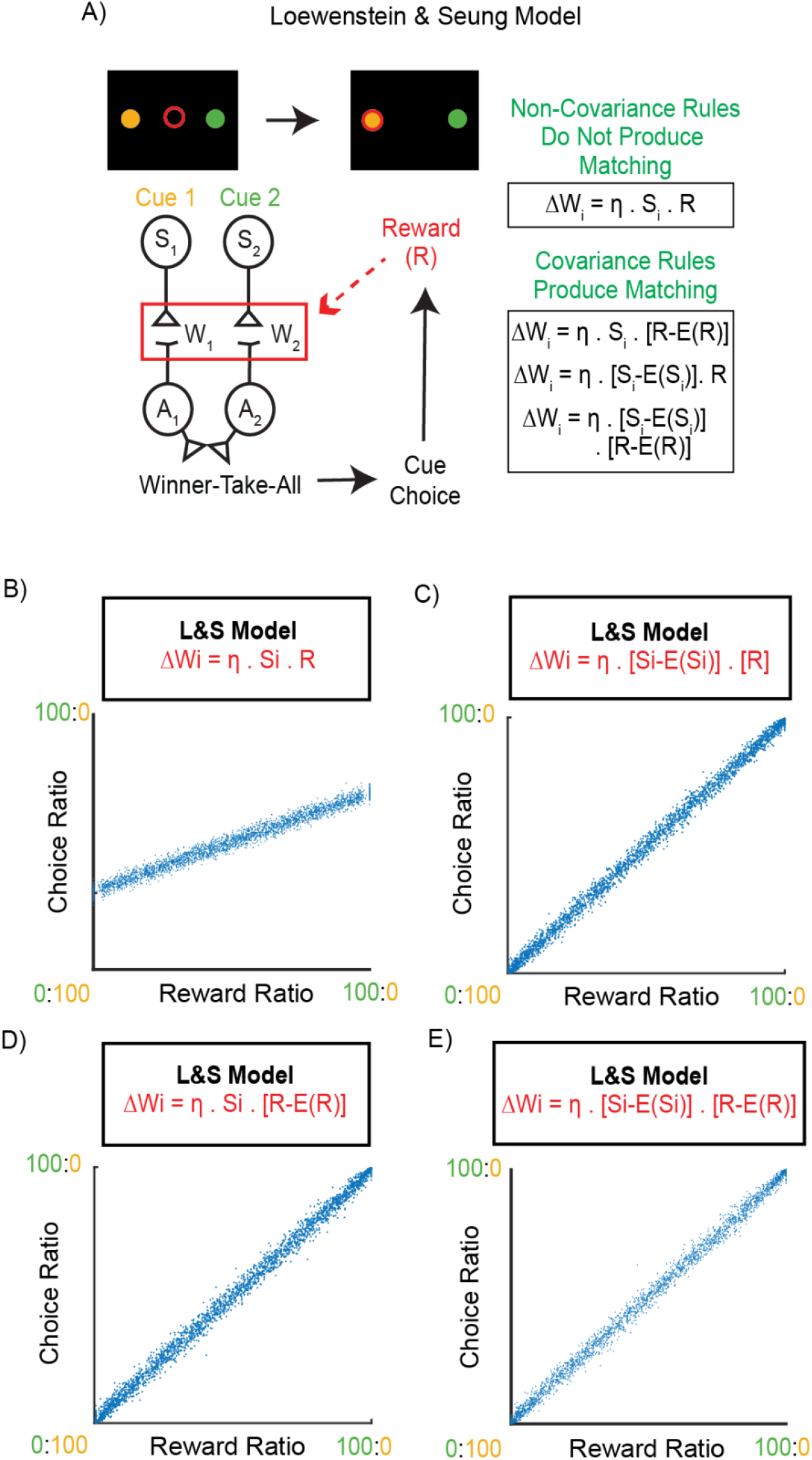
Covariance-based learning rules are necessary for operant matching. **A** Description of the model developed by Loewenstein and Seung to study the requirements of matching behavior. The neural network consists of sensory neurons S_1_ and S_2_ that respond to one of the two simultaneously provided stimuli and synapse onto motor neurons A_1_ and A_2_ via synapses with weights W_1_ and W_2_. Choices are determined via a winner-take-all computation downstream of motor neurons. Upon choice, weights are updated according to one of the shown plasticity rules (*boxes-right*). Here, S_i_ is the activity of the i^th^ sensory neuron; R represents the presence or absence of reward; E(S_i_) is the mean or expectation of the activity of S_i_ and E(R) is the expectation of reward. **B** Block-averaged choice ratios versus reward ratios (n = 2000 simulations) from data simulated using a non-covariance in the model described in 4A are plotted against each other. **C** Block-averaged choice ratios versus reward ratios (n = 2000 simulations) from data simulated using a covariance-based rule [S_i_-E(S_i_)]*R in the model described in 4C are plotted against each other. **D** Block-averaged choice ratios versus reward ratios (n = 2000 simulations) from data simulated using a covariance-based rule S_i_*[R-E(R)] in the model described in 4A are plotted against each other. **E** Block-averaged choice ratios versus reward ratios (n = 2000 simulations) from data simulated using a covariance-based rule [S_i_-E(S_i_)]*[R-E(R)] in the model described in 4A are plotted against each other.

**Supplementary Figure 5:**
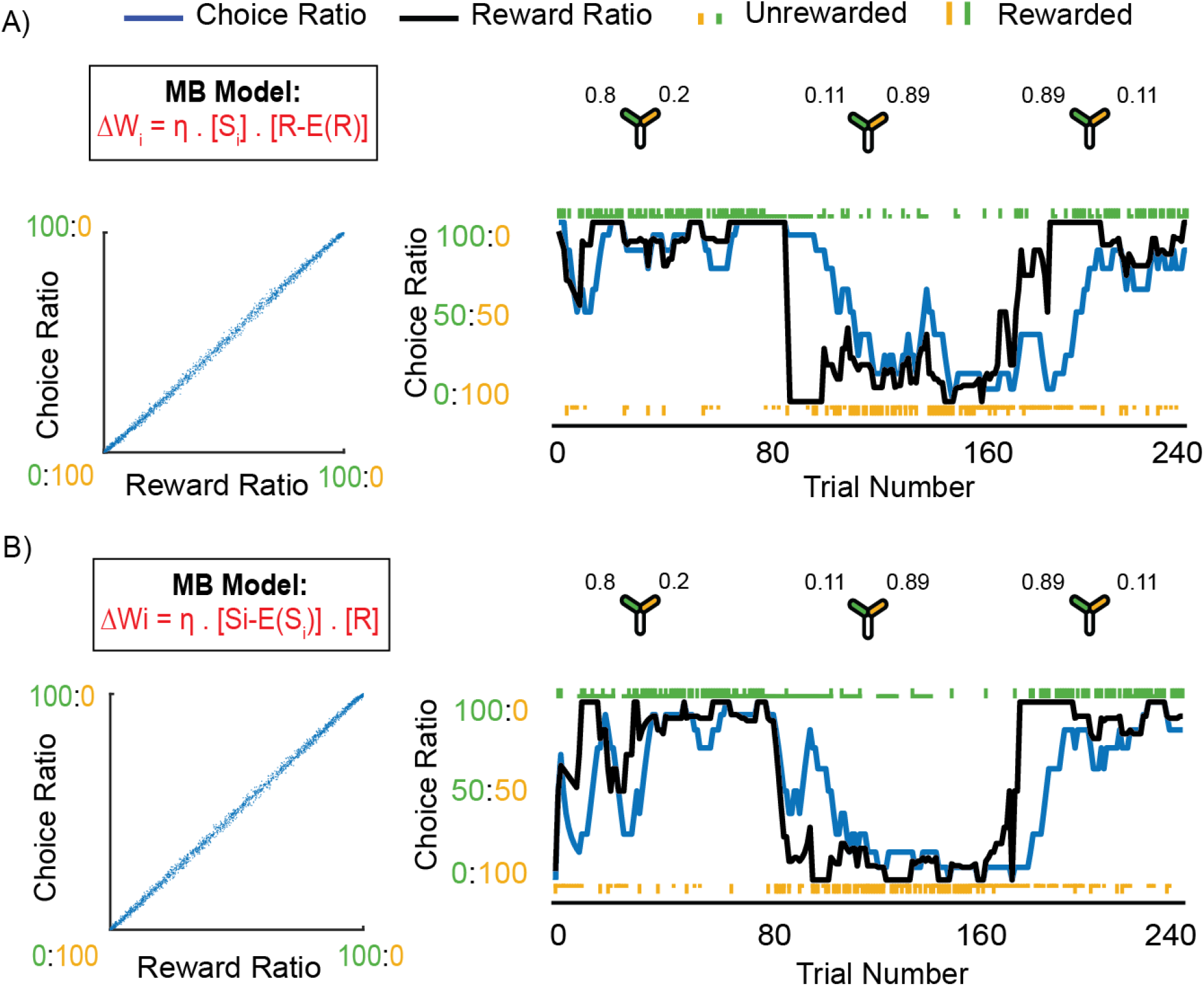
Models using covariance-based learning rules produce behavior more similar to real fly behavior. **A** *Left:* Block-averaged choice ratio produced by the S_i_*[R-E(R)] covariances-based rule (*box*) plotted against reward ratio. The model exhibits matching behavior (slope is 1). *Right:* An example simulation showing the performance in a 3 block task of a model incorporating a covariance-based rule S_i_*[R-E(R)]. Task reward contingencies are the same as shown for the example fly in Fig. 2A. **B** Same as (A), but simulated with a [S_i_-E(S_i_)]*R rule. *Left:* The model exhibits matching behavior (slope is 1). *Right:* performance in a 3 block task where reward contingencies are the same as shown for the example fly in Fig. 2A.

**Supplementary Figure 6:**
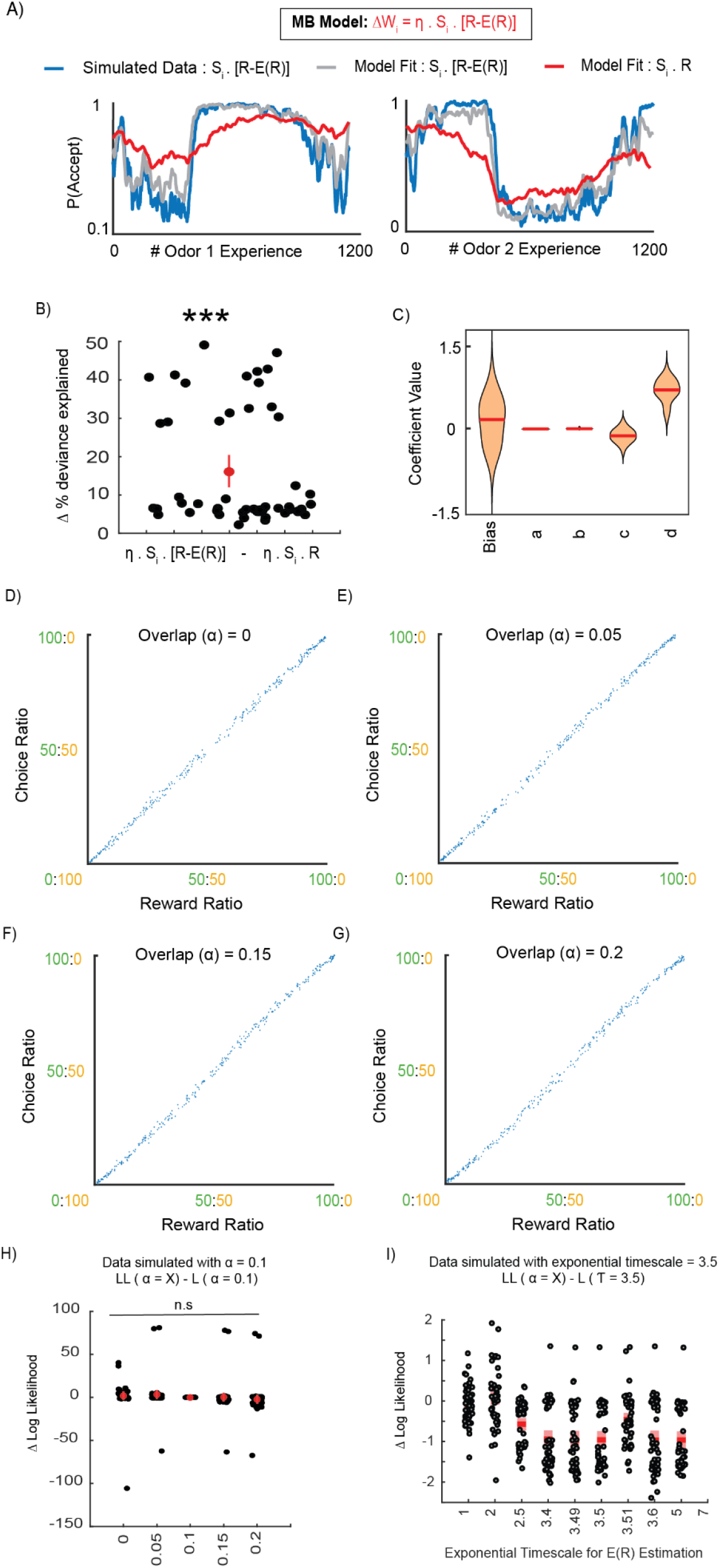
Extent of sensory overlap does not affect the behavior of our model. **A** Example simulated data showing the probability of accepting odor 1 *(left)* and odor 2 *(right)* (blue), simulated using a covariance-based rule with reward expectation, and fit using an MB-inspired regression model (A) that incorporates either the same rule (gray), or a non-covariance rule (red). The predictions resulting from the model using the covariance-based rule is a better fit for the simulated data. **B** Change in percentage deviance explained, computed by subtracting the percentage deviance explained of the non-covariance-based regression model from the covariance-based rule, plotted for each simulation (n = 50). The covariance-based rule better fits the simulated behavior than the non covariance-based rule (Wilcoxon signed-rank test: p = 6.7595 * 10^−9^) **C** Regression coefficients assigned to each term of the learning rule when the MB-inspired regression model using a covariance-based rule with reward expectation was fit to the simulated behavior. Note the non-zero weight of the c and d terms. **D** Block-averaged choice ratios versus reward ratios (n = 300 simulations) from data simulated using a covariance-based rule with reward expectation are plotted against each other. Sensory overlap α = 0. **E** Block-averaged choice ratios versus reward ratios (n = 300 simulations) from data simulated using a covariance-based rule with reward expectation are plotted against each other. Sensory overlap α = 0.05. **F** Block-averaged choice ratios versus reward ratios (n = 300 simulations) from data simulated using a covariance-based rule with reward expectation are plotted against each other. Sensory overlap α = 0.15. **G** Block-averaged choice ratios versus reward ratios (n = 300 simulations) from data simulated using a covariance-based rule with reward expectation are plotted against each other. Sensory overlap α = 0.2. **H** Change in percentage deviance explained, computed by subtracting the percentage deviance explained of the covariance-based regression model with overlap = 0.1, from that of regression models with covariance-based rules and different amounts of overlap (0,0.05,0.15,0.2). Points are plotted for each simulation (n = 50). Simulations were run with overlap α = 0.1. **I** Change in percentage deviance explained, computed by subtracting the percentage deviance explained of the covariance-based regression model with exponential timescale (τ)= 3.5, from that of regression models with covariance-based rules and different exponential timescales. Points are plotted for each simulation (n = 50). Simulations were run with exponential timescale (τ)= 3.5.

**Supplementary Figure 7:**
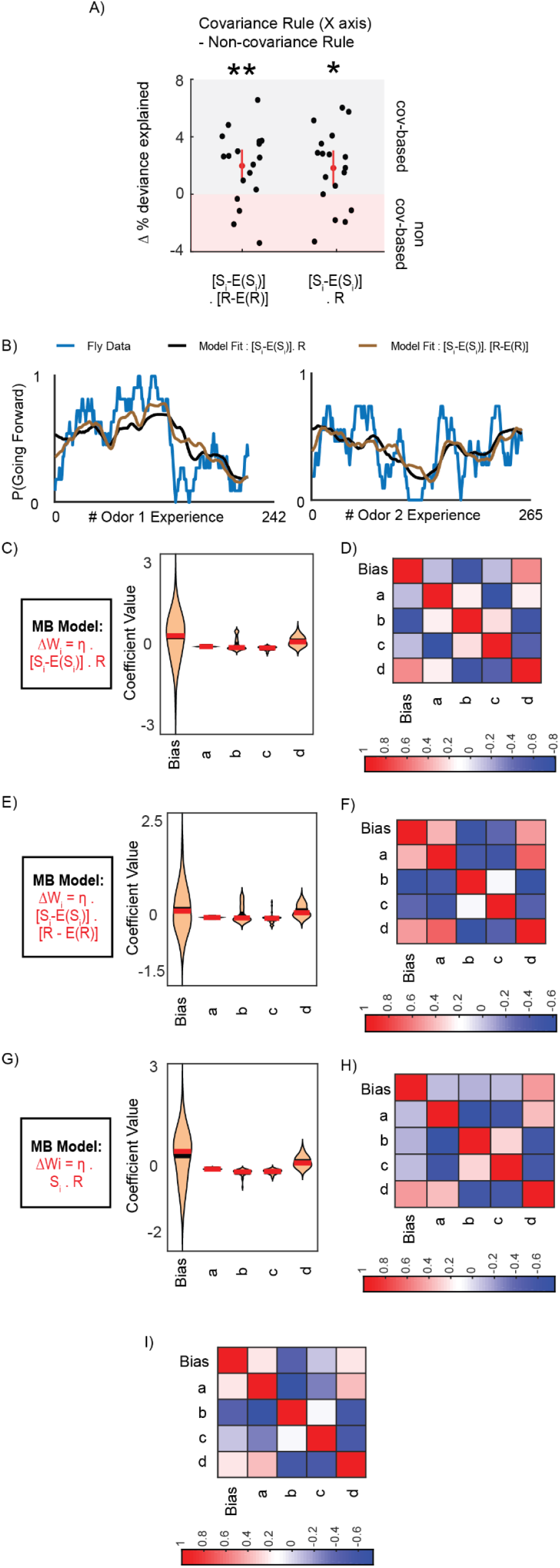
Covariance-based learning rules are better predictors on individual choice behavior. **A** Change in percentage deviance explained, computed by subtracting the percentage deviance explained of the non-covariance-based model from two models with covariance-based rules (*left:* incorporating both stimulus and reward expectations; *right:* incorporating just stimulus expectation) plotted for each fly (n = 18). Covariance-based rules were more predictive of fly behavior on average (Wilcoxon signed-rank test:*left*: p=0.0074; *right*: p = 0.0168). **B** Example fly data showing probability of accepting odors as a function of odor experience number for odor 1 *(left)* and odor 2 *(right)*(blue) fit using an MB-inspired regression model (Fig. 5A) that incorporates a covariance rule with either sensory and reward expectations (brown), or just sensory expectations (black). **C** Regression coefficients assigned to each term of the learning rule when the MB-inspired regression model using a covariance rule with sensory expectation was fit to the flies’ behavior. **D** Correlation between regression coefficients resulting from MB-inspired regression model using a covariance rule with sensory expectation. **E** Regression coefficients assigned to each term of the learning rule when the MB-inspired regression model using a covariance rule with sensory and reward expectations was fit to the flies’ behavior. **F** Correlation between regression coefficients resulting from MB-inspired regression model using a covariance rule with sensory and reward expectations. **G** Regression coefficients assigned to each term of the learning rule when the MB-inspired regression model using a non-covariance rule was fit to the flies’ behavior. **H** Correlation between regression coefficients resulting from MB-inspired regression model using a non-covariance rule. **I** Correlation between regression coefficients resulting from MB-inspired regression model using a covariance rule with reward expectation.

**Supplementary Figure 8:**
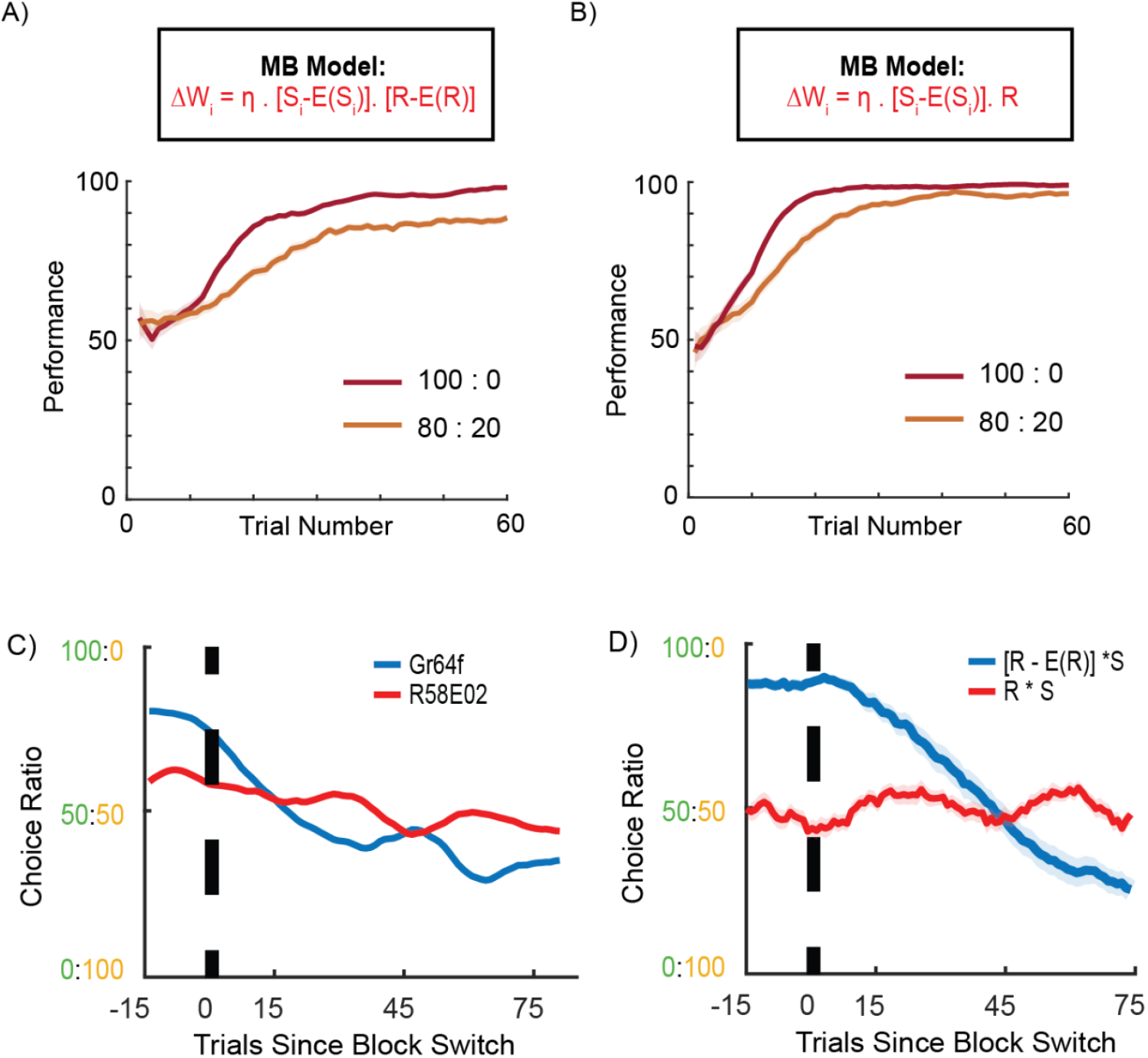
Covariance-based learning rules produce similar behavior in 100:0 and 80:20 tasks. **A** Simulated instantaneous performance plotted as a function of trial number (defined as the percentage of choices towards the option with higher pre-defined reward probability in a 10 trial window) of an agent using a covariance-based rule with sensory and reward expectations in 80:20 (orange) and 100:0 (red) reward conditions. **B** Simulated instantaneous performance plotted as a function of trial number (defined as the percentage of choices towards the option with higher pre-defined reward probability in a 10 trial window) of an agent using a covariance-based rule with sensory expectation in 80:20 (orange) and 100:0 (red) reward conditions. **C** Change in instantaneous choice ratio around block changes. Flies trained with Gr64f activation in blue, DAN activation in red **D** Change in instantaneous choice ratio around block changes from simulated data. Agents with a covariance-based rule with reward expectation in blue. Agents with non-covariance rule in red.

**Supplementary Figure 9:**
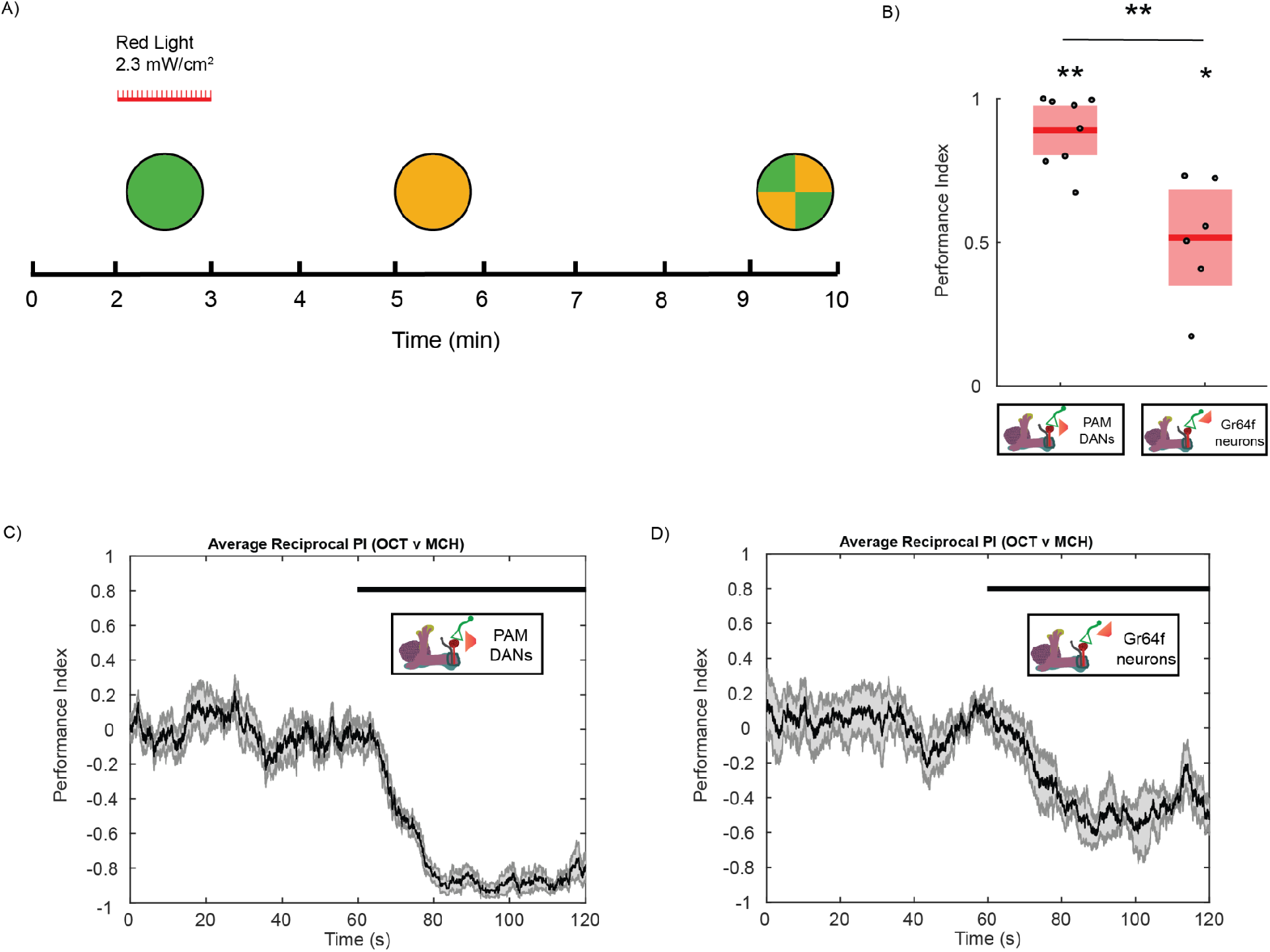
Circular arena experiments to control for rewarding red LED intensity. **A** Schematic of the experimental paradigm used to train flies in the circular arena. LED intensity chosen to be 2.3 mW/cm^2^ to match intensity in the Y-arena. **B** Time averaged performance index plotted for DAN trained and Gr64f trained flies show that both learn to prefer the reward-paired odor (Wilcoxon signed-rank test: Gr64f - p = 0.0312; DAN - p = 0.0078). **C** Performance index plotted over timecourse of testing period (minute 9-10 in (A)) for DAN trained flies. **D** Performance index plotted over timecourse of testing period (minute 9-10 in (A)) for Gr64f trained flies.

**Supplementary Information 1:**
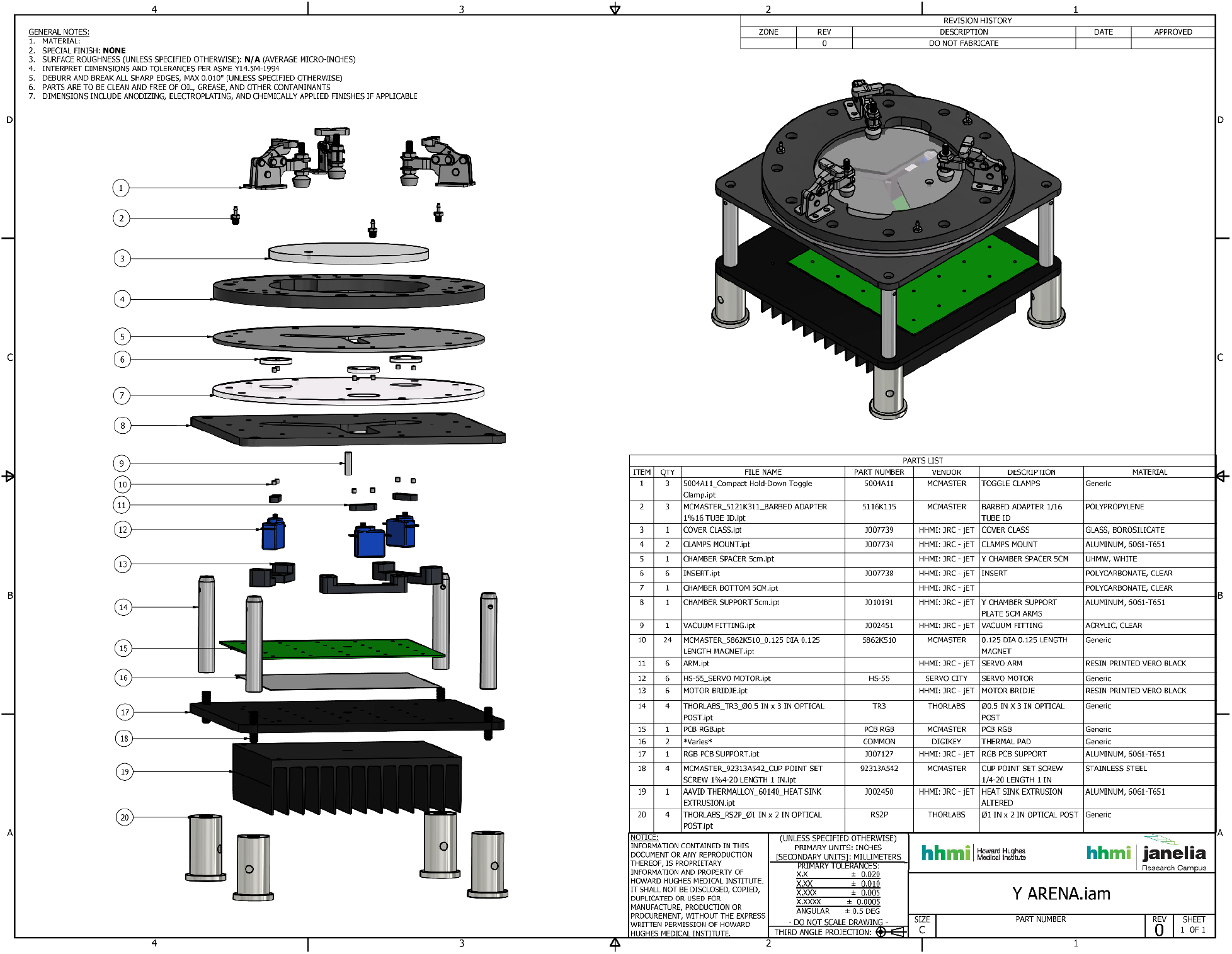
Description of the Y-arena.

